# Molecular characterization of the stress network in the human brain

**DOI:** 10.1101/661587

**Authors:** Mandy Meijer, Arlin Keo, Judith M.C. van Leeuwen, Oleh Dzyubachyk, Onno C. Meijer, Christiaan H. Vinkers, Ahmed Mahfouz

## Abstract

The biological mechanisms underlying inter-individual differences in human stress reactivity remain poorly understood. We aimed to identify the molecular underpinning of neural stress sensitivity. Linking mRNA expression data from the Allen Human Brain Atlas to task-based fMRI revealed 201 differentially expressed genes in cortex-specific brain regions differentially activated by stress in individuals with low or high stress sensitivity. These genes are associated with stress-related psychiatric disorders (e.g. schizophrenia and anxiety) and include markers for specific neuronal populations (e.g. *ADCYAP1, GABRB1, SSTR1*, and *TNFRSF12A*), neurotransmitter receptors (e.g. *GRIN3A, SSTR1, GABRB1*, and *HTR1E*), and signaling factors that interact with the corticosteroid receptor and hypothalamic-pituitary-adrenal axis (e.g. ADCYAP1, *IGSF11, and PKIA*). Overall, the identified genes potentially underlie altered stress reactivity in individuals at risk for psychiatric disorders and play a role in mounting an adaptive stress response, making them potentially druggable targets for stress-related diseases.

## 1. INTRODUCTION

Stress is a major risk factor for the development of a wide range of psychiatric disorders, including schizophrenia and depression.^1^ Inter-individual differences in how the brain responds to stress depend on intrinsic (*e.g.* genetic and developmental) as well as on extrinsic (*e.g.* hormonal) factors.^2^ The neural correlates underlying stress reactivity are currently a growing topic of investigation.^3–5^ In healthy individuals, acute stress causes a shift in neural networks by suppressing the executive control network and activating the salience network and default mode network (DMN).^6^ One hypothesis is that stress vulnerability is the result of maladaptive changes in the dynamic response of these neural networks, either during the acute phase, during the recovery period in the aftermath of stress, or both.^2^ Moreover, acute social stress deactivates the DMN in the aftermath of stress during emotion processing in healthy controls but not in siblings of schizophrenia patients who are at-risk for several psychiatric disorders.^7, 8^ Yet, the molecular mechanisms underlying differences in brain reactivity to stress in humans remain unknown as access to the tissue of interest in humans is limited.

Nevertheless, stress-related brain regions and networks as identified by fMRI can be further characterized based on transcriptomic signatures. Mapping gene expression atlases of the healthy brain to imaging data allows the identification of the molecular mechanisms underlying imaging phenotypes. Previous studies have identified gene expression patterns associated with structural brain changes in autism spectrum disorders, Huntington’s disease and the onset of schizophrenia.^9–12^ Similarly, mapping resting-state fMRI and connectivity data onto gene expression atlases has led to identification of molecular profiles underlying these fMRI networks.^13–15^

In this study, we examined the putative molecular signatures of brain regions linked to stress reactivity. We linked gene expression data from the Allen Human Brain Atlas (AHBA) to an fMRI-stress network (**Figure 1**). In short, we found that the stress network was enriched for genes associated to specific subtypes of neurons (*i.e.* components of the cortical circuitry) with genetic relevance for psychiatric disorders, and for signaling factors and proteins that interact with the activation of the Hypothalamic-Pituitary-Adrenal axis (HPA-axis) and response to glucocorticoids. These all constitute potential targets for directed pharmacotherapy in stress related disorders.

**Figure 1.**
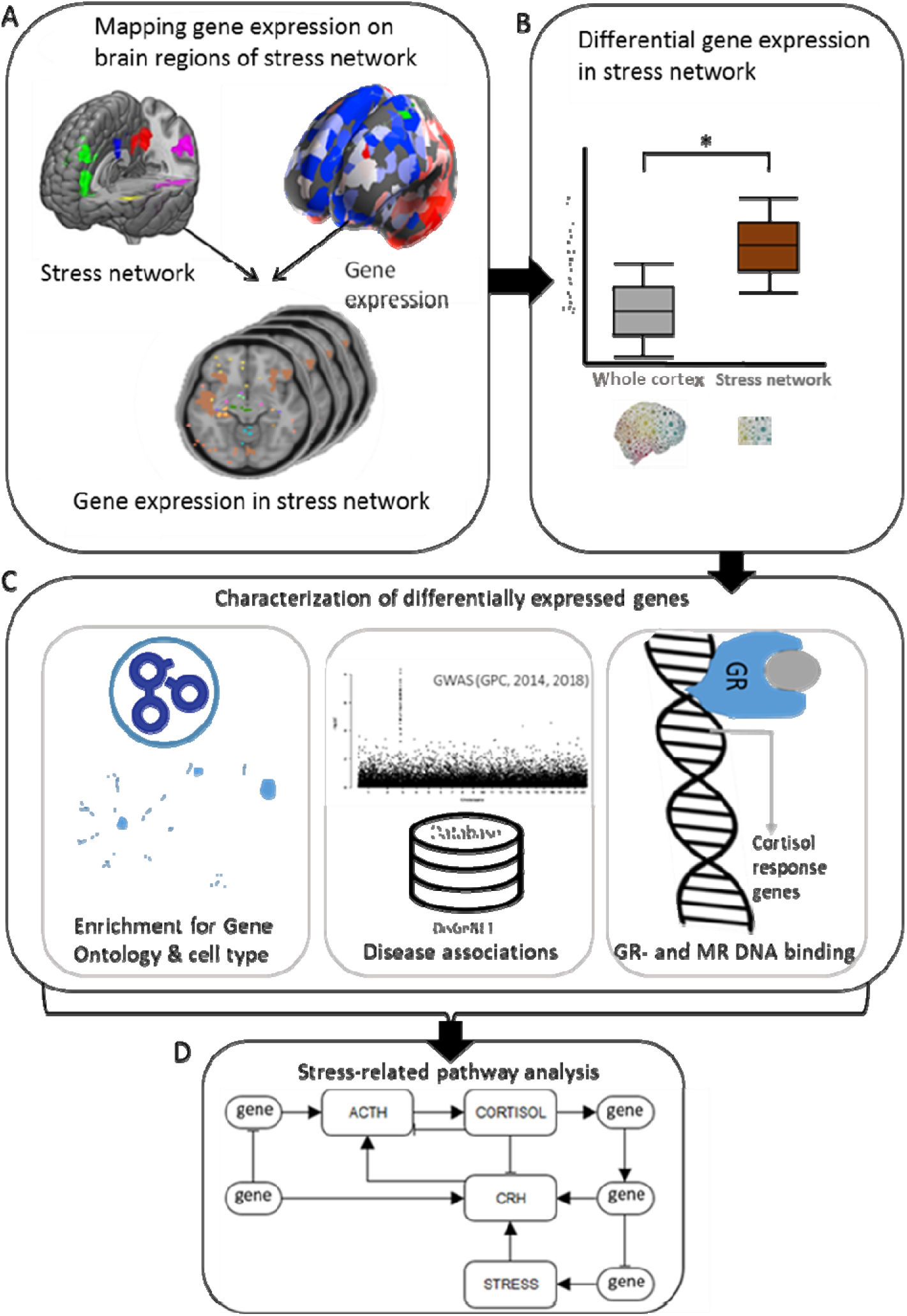
Study overview. (A) Cortical brain regions vulnerable to stress (= stress network) during an emotion processing task were assessed in an fMRI study. All brain regions showed higher stress-induced brain activity following an acute social stressor in at risk individuals (healthy siblings of schizophrenia patients). The fMRI data was mapped to the AHBA resulting in an overlay of the fMRI and gene expression data. (B) With this overlay, differential gene expression between the brain regions vulnerable to stress and the rest of the cortex were assessed. (C) Differentially expressed genes were consequently characterized by identifying enrichment for gene ontology and cell type markers, associations with stress-related diseases and enrichment for cortisol responsive genes. (D) Information provided by the previous analyses was used to build a model of a molecular pathway underlying human stress reactivity.

## 2. METHODS

### 2.1 Defining the stress network

Based on a previous study, we selected brain regions that were differentially affected by stress in individuals with high and low stress sensitivity.^6^ In this study, there were four experimental groups: control-no-stress (n=19), control-stress (n=20), sibling-no-stress (n=20) and sibling-stress (n=19) (Table 1). Before scanning, participants in the stress groups underwent a Trier Social Stress Test^16^ and 30 minutes after the onset of the test, participants performed an emotion-processing task in the magnetic resonance imaging (MRI) scanner based on the International Affective Picture System^17^ during which pictures were presented that had to be rated as either neutral, positive or negative. All participants in this experiment gave written informed consent and the experiment was approved by the Utrecht Medical Center ethical review board and performed according to the guidelines for Good Clinical Practice and the declaration of Helsinki. Based on a 2×2 ANOVA (control/sibling × stress/no-stress) voxel-wise analysis, several brain regions that responded differently to all pictures after acute social stress in siblings compared to healthy individuals were identified. These regions include key nodes of the DMN (posterior cingulated cortex/precuneus and medial prefrontal cortex) and salience network (anterior insula), as well as the superior temporal gyrus, middle temporal gyrus, middle cingulate gyrus, ventrolateral prefrontal cortex, precentral gyrus and cerebellar vermis (**Figure 2A**). We selected and present in the figures the cortex-specific brain regions for the initial analyses to prevent that our results are being driven by differences between the cortex and subcortex. Analyses on all brain regions in the stress network can be found in the supplementary text.

**Table 1.** Group characteristics of fMRI study.

|  | **Con-no-stress** | **Con-stress** | **Sib-no-stress** | **Sib-stress** | **p-value** |
| --- | --- | --- | --- | --- | --- |
| **N** | 19 | 20 | 20 | 19 |  |
| **Age (years)** | 32.6 (8.5) | 34.8 (9.1) | 33.8 (10.8) | 32.5 (7.4) | 0.836<sup>a</sup> |
| **Handedness (% right)** | 89.5 | 95 | 70 | 89.5 | 0.194<sup>b</sup> |
| **Educational level** | 7.6 (2.7) | 7.1 (1.9) | 7.0 (1.6) | 7.4 (1.5) | 0.688<sup>a</sup> |
| **Body Mass Index** | 24.1 (2.7) | 24.2 (2.1) | 24.0 (3.0) | 24.9 (3.9) | 0.774<sup>a</sup> |
| **Ethnicity (% Caucasian)** | 84.2 | 90 | 90 | 84.2 | 0.900<sup>b</sup> |
| **Smoker (% yes)** | 5.3 | 35 | 30 | 31.6 | 0.132<sup>b</sup> |

Con = control; sib = sibling of schizophrenia patients.
Mean values (SD) are denoted for age, education, and body mass index. All other values are reported in frequency.
a = one-way-ANOVA
b = chi square test

**Figure 2.**
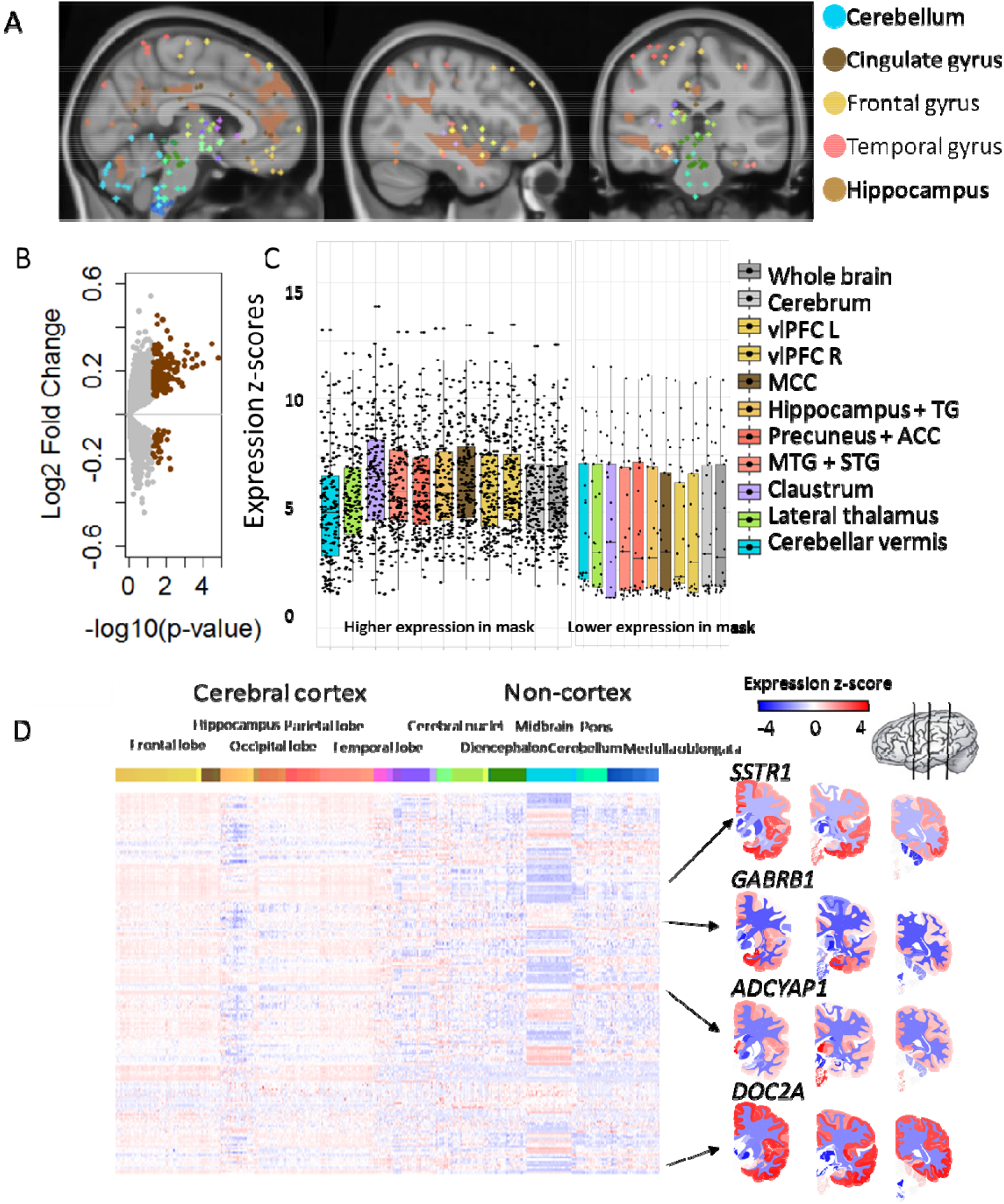
Differentially expressed genes in brain regions vulnerable to stress can be identified using gene expression atlases. (A) Brain regions in the stress network are present throughout the brain (including cerebellum, cingulate gyrus, frontal gyrus, temporal gyrus, and hippocampal formation). For the analysis, all regions in the cortex were combined. (B) Differential gene expression was determined for the cortical stress network compared to the rest of the cortex. Significant genes (BH-adjusted p-value < 0.05 have higher expression in the stress network. Grey dots represent non-significant genes and brown dots represent significant genes based on meta-analysis across all six AHBA donors. (C) The box plots show the expression of the higher (left) and lower (right) expressed genes compared to the rest of the cortex in the brain regions of interest from the stress network. (D) In the whole brain, differentially expressed genes show mostly high expression levels in the cortex and low expression levels in non-cortical brain regions. In the heatmap, each row represents a gene and each column represents a sample and all samples of the AHBA are illustrated here. On the right, coronal brain sections for the genes *SSTR1, GABRB1, ADCYAP1,* and *DOC2A* are presented. Colors indicates high (red) and low (blue) expression levels.

### 2.2 Allen Human Brain Atlas (AHBA)

Gene expression data from six healthy brains were acquired from the AHBA.^18^ In this microarray dataset, probes were mapped to genes as previously described.^19^ Z-scores for normalized gene expression levels from the AHBA were calculated separately for each of the six individual brains. Gene expression data were linked to an fMRI-based stress network according to the MNI coordinate system, such that samples of the AHBA exactly overlap with the corresponding fMRI voxels. For all samples in the AHBA, we determined whether they were located in the cortical stress network for all six donors separately. The gene expression levels of the AHBA samples were extracted and resulted in expression data of 19,992 genes in 111 and 1839 brain samples in- and outside the cortical stress network, respectively.

### 2.3 Differential gene expression in the cortical stress network

To identify genes differentially expressed between the cortical stress network and the rest of the cortex, we analyzed each of the six brain donors separately. Differential expression was determined for the cortical stress network altogether as one mask. For each gene, we combined effect sizes (difference in mean expression between the brain stress network and the rest of the brain) across donors using a meta-analysis approach from the ‘metafor’ 2.0-0 R package. In brief, a random effects model was used, taking into account the within-brain and between-brain variance, which was estimated with the Dersimonian-Laird model. Variances and confidence intervals needed for the meta-analysis were calculated using the escalc-function. Genes were considered to be differentially expressed at an BH-adjusted p-value < 0.05 (Benjamini-Hochberg (BH) correction).

We also performed analysis on the whole brain (differentially expressed genes BH-adjusted p-value < 0.05 and log_2_(Fold Change) > |1|). Given the large difference in the transcriptional profile of the cerebellum compared to the rest of the brain^20^, we excluded the cerebellum from the whole brain analysis. In addition, we performed the differential expression analysis between samples inside and outside the stress network for each of the following brain regions separately: cerebral cortex (Cx), frontal gyrus (FG), cingulate gyrus (CgG), cerebellum (Cb), and the hippocampal formation (HiF). Other anatomical regions contained too few samples (< 2 in the mask) to perform the analysis on these particular structures separately.

We used a bootstrapping approach to assess the robustness of our results with respect to the imbalance between the number of AHBA samples inside and outside the cortical stress network (111 inside and 1839 outside). We randomly selected 111 samples from the whole cortex, regardless of their location inside or outside the stress network and compared gene expression profiles of these brain samples with the original set of 111 samples inside the cortical stress network. We repeated this process 1000 times to assess the reproducibility of the differentially expressed genes

### 2.4 Gene Ontology (GO) enrichment analysis

To characterize the functionality of the differentially expressed genes, a GO enrichment analysis was performed. The list of unranked differentially expressed genes was uploaded to GOrilla (Gene Ontology Enrichment Analysis and Visualization Tool).^21^ As a background list, the top 20% of genes with the highest expression level in the cortex was used, to correct for non-selective ontologies. GO terms were considered significant when the p-values < 0.001 (Fisher’s exact test) after BH-correction.

### 2.5 Cell type enrichment analysis

We assessed whether the differentially expressed genes were enriched for cell type markers.^22^ Genes with a 20-fold higher expression in neurons (628 marker genes), oligodendrocytes (186 marker genes), astrocytes (332 marker genes), microglia (520 marker genes) and endothelial cells (456 marker genes) were considered to be markers for that cell type. Since most of our AHBA samples were located inside the cortex, we used a set of brain-region-specific markers and focused on 18 cortical cell types.^23^ Details on markers can be found on https://pavlab.msl.ubc.ca/data-and-supplementary-information/supplement-to-mancarci-et-al-neuroexpresso. Finally, to assess which neuronal cell types might be involved in stress sensitivity, single cell RNA sequencing data of the middle temporal gyrus of the human neocortex from the Allen Brain Institute^24^ (http://celltypes.brain-map.org/rnaseq/human) were used. The sum of the log_10_ values of the counts per differentially expressed gene were calculated for each cell cluster separately.

### 2.6 Enrichment analysis of disease-associated genes

To assess whether the differentially expressed genes are associated to stress-related psychiatric disorders, a disease-associated gene enrichment analysis was performed based on existing Genome-Wide Association Studies (GWAS) including schizophrenia^25,26^, Bipolar Disorder^31^, and Major Depressive Disorder^32^, and stress-related diseases such as Post-Traumatic Stress Disorder, as well as non-stress-related diseases (e.g. Huntington and osteoporosis) based on disease gene sets from DisGeNET^27^. As non-disease control conditions, genes associated to height and waste-hip ratio were included in the analyis.^28, 29^ The schizophrenia, MDD and BP GWAS loci were considered to be associated if they reached genome-wide significance of p < 5*10^−8^. Intersections of loci based on GENCODE with UCSC hg19/NCBI build 37 position were used to map loci to risk genes by the authors of the GWAS.^25, 30, 31^ These annotations were used for the enrichment analyses. All genes assessed in the AHBA that were not associated to a disease or trait were used as background test in the Fisher-test. FDR-correction was applied over the amount of enrichment tests.

To assess the enrichment of disease-related gene sets in intercellular signaling genes, neuropeptides and receptor genes were selected from the differentially expressed genes. Odds ratios (ORs) were calculated for the set of neuropeptides and receptors for each disease as a measurement of effect size, (i.e. the increased chance of a peptide or receptor being present in the set of differentially expressed genes). For this, the number of receptors found within the trait was compared to all the receptors that were measured in the AHBA (1203 receptors), based on the gene annotation of the AHBA. Gene names that included the word ‘receptor’ were selected and this list was manually verified whether the gene was a receptor or a modulator. The ORs for the neuropeptides were calculated in the same way, based on a list of neuropeptides available from NeuroPep.^32^

### 2.7 Mineralocorticoid and glucocorticoid DNA binding loci

MR and GR DNA binding loci under stress in the rat hippocampus were previously assessed.^33^ We identified sets of genes with GR-specific, MR-specific and GR-MR-overlapping DNA binding loci, i.e. potential target genes. To predict glucocorticoid sensitivity of our differentially expressed genes, we assessed whether these sets of targets were enriched among the differentially expressed genes.

### 2.8 Enrichment statistics for GO, cell type, disease-associated genes and receptor binding

Enrichments were assessed based on Fisher’s Exact Tests and odds ratios (ORs) were calculated as a measurement of effect size for the enrichments. An OR of 1 indicates no effects, whereas an OR > 1 and 0 < OR < 1 reflects enrichment and depletion, respectively. All p-values were corrected for multiple testing using Benjamini-Hochberg method and a BH corrected p-value < 0.05 was considered to be significant, unless stated otherwise.

## 3. RESULTS

### 3.1 Differentially expressed genes in the stress network

We identified the gene expression signatures of the cortical stress network with altered stress-induced activity by determining which genes are differentially expressed in the stress network compared to the rest of the cortex. Using a meta-analysis approach to combine results across all donors of the AHBA (n=6), we identified 201 differentially expressed genes (BH-adjusted p < 0.05, **Figure 2B; Table S1**). Among those genes, 177 were higher expressed, while the other 24 genes were lower expressed in the cortical stress network compared to the rest of the cortex. Using a bootstrapping approach (see 2.3), we found the identified set of genes to be highly robust to the imbalance between the number of AHBA samples inside and outside the stress network (in 83% of our 1000 iterations, only genes out of our initial 201 differentially expressed gene list were found).

We also identified the gene expression signatures of the stress network with altered stress-induced activity by determining which genes are differentially expressed in the stress network compared to the rest of the brain minus the cerebellum. Using the same meta-analysis approach, we identified 261 differentially expressed genes (BH-adjusted p < 0.05 and log_2_(fold-change) > |1|). A full description of the results, including tables and figures, can be found in the **supplementary text**. However, due to the higher representation of cerebral cortex samples in the brain regions vulnerable to stress (109 out of 127; 91%) compared to the rest of the brain minus the cerebellum (1,950 out of 3,225; 60%), differentially expressed genes in the whole brain stress network were also differentially expressed between cortical and non-cortical samples (222 out of 261 genes were also differentially expressed in the top 10% difference between cortex and non-cortex, p < 0.00001). Therefore, we chose to focus on the cortex-specific stress network.

The differentially expressed genes in the cortical stress network generally showed high expression values in the cortex but not the hippocampus, and mostly low expression levels in non-cortical areas (**Figure 2D**). The two most differentially expressed genes in the stress-specific cortical regions are Tumor necrosis factor receptor superfamily member 12A (*TNFRSF12A*) (BH-adjusted p-value = 0.006, log_2_(FC) = −0.24) and Exosome Component 6 Pseudogene (*LOC392145*) (BH-adjusted p-value = 0.009, log_2_(FC) = 0.38). *TNFRSF12A* may module cellular adhesion to matrix proteins, whereas *LOC392145* is an mRNA transport regulator.

### 3.2 Functionality and cell-type specificity of differentially expressed genes in the stress network

A GO term enrichment analysis was performed to assess whether the differentially expressed genes in the stress network are enriched for specific functions. The differentially expressed genes were enriched for GO terms involved in neuronal development and neurogenesis, synaptic signal transmission, and glutamate receptor signaling (**Figure 3A and Table S1**). Genes involved in most processes based on GO terms (at least assigned to five out of ten GO terms) include *SHANK, GRIN3A, CNTN4* and *ADCYAP1*. Enrichment analysis for cellular components indicated that the proteins coded by the differentially expressed genes were mainly found at the synapse, reflecting both the high expression of the genes in the synapse-dense cerebral cortex, and a potential role for synaptic proteins as determinants for the differential activation.

**Figure 3.**
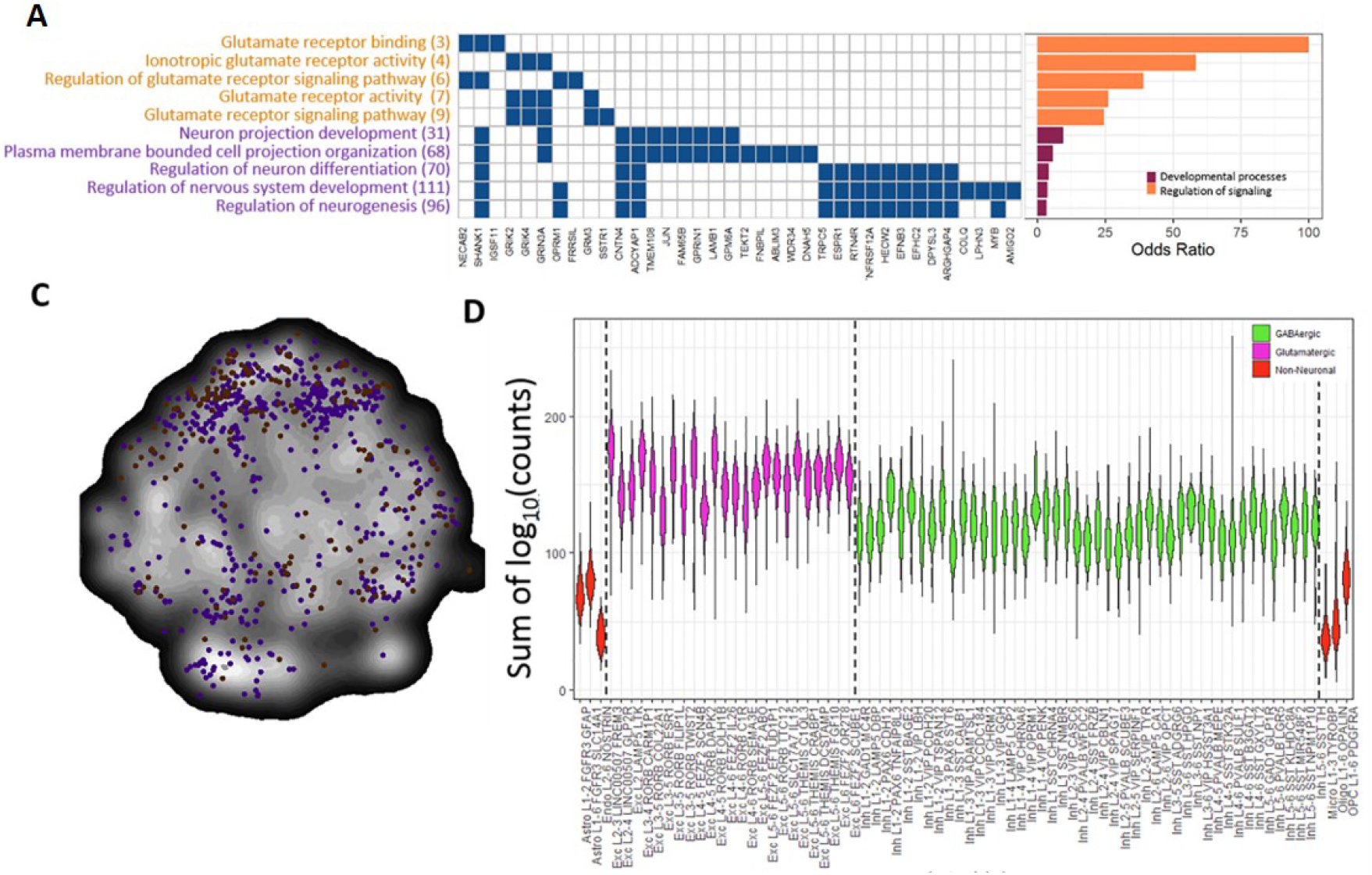
Functionality of differentially expressed genes in the stress network. (A) Differentially expressed genes annotated to one of the GO terms were assigned to multiple GO terms and thus involved in multiple processes. Between parenthesis, the total number of genes assigned to the GO term is depicted. On the right side of the graph, ORs are displayed per GO term. (B) Differentially expressed genes (brown), neuronal marker genes (purple) and overlapping genes (yellow) are plotted in a t-SNE plot generated using BrainScope.nl^20^, where points close together represent genes with similar gene expression profiles. The differentially expressed genes show a similar profile in the t-SNE plot as neuronal cell markers (purple). (C) ORs for different neuronal subtypes. ORs were considered significant when the BH-adjusted p-value < 0.05 (*). Neuronal subtypes without a marker represented in the differentially expressed genes are not illustrated in the graph. (D) The sum of the log10 values of the counts per gene is plotted for each cell cluster. Green clusters belong to GABAergic cells, purple clusters to glutamatergic cells and red clusters to non-neuronal cells.

Next, we identified the specific cell types underlying the differential gene expression levels in the cortical stress network using enrichment analysis of cortical cell-type markers.^22^ Enrichment was found for neuronal cell markers (BH-adjusted p = 5*10^−5^), including *ADCYAP1, DPYSL3, INSM1, PKIA, SSTR1, NOL4, BAIAP3, KCNB2, FAM65B, ABLIM3, TEKT2, SHANK1, DACT1, PCBP3, SCN3B, LMO3, CA10, LRRTM4, SYT16, GPRIN1, TMEM200A, LRRC3B, GRIN3A,* and *PNCK*. However, no specific subtype of neuron was in particular enriched. Enrichment was also found for astrocytes (*ATP2B4, PTCH1, FABP7, IGSF11, KCNN3, GRM3, GABRB1, PTX3,* BH-adjusted p = 0.024). The list of differentially expressed genes included a few microglia (*TMEM52, F13A1, MSH5, ARHGAP4, NOD2,* and *TNFRSF12A*), endothelial cell (*LAMA1, LAMB1, C1orf115, DOK4,* and *MICB*), and oligodendrocyte markers (*EFNB3* and *TYRO3*), although not significantly enriched (BH-adjusted p = 0.924). Moreover, we found that neuronal markers showed a partially overlapping distribution in a t-Distributed Stochastic Neighbor Embedding (t-SNE) map of all genes across the whole brain as the differentially expressed genes in, indicating that neuronal markers and the differentially expressed genes show the same expression patterns across cortical areas (**Figure 3B**) and thus differential activity may depend on neuronal gene expression.

Using a human-specific single cell RNA-sequencing data of the medio-temporal gyrus^24^, we found the differentially expressed genes to be mainly enriched in glutamatergic excitatory neurons compared to GABAergic and non-neuronal cells, using a Wilcoxon rank test (p-value = 2.2*10^−^16, **Figure 3D**).

### 3.3 Differentially expressed genes in stress network are associated to stress-related diseases

We hypothesized that the differentially expressed genes in the stress network would be associated to the genetic background of psychiatric disorders, particularly for stress-related brain disorders, as stress plays a major role in the development of these disorders. Using genetic variants from GWAS of the Genomics of Psychiatry Consortium^25, 26^, we assessed whether schizophrenia-associated risk loci are enriched in the set of differentially expressed genes. Indeed, schizophrenia risk genes were enriched in the differentially expressed genes in the stress network (Fisher Exact test, BH-adjusted p-value = 0.015). The schizophrenia risk genes *CNTN4, GRM3, FUT9, SATB2, GPM6A, COQ10B, DOC2A,* and *NISCH* were present in our differentially expressed genes, and all except one (COQ10B) were higher expressed in the cortical brain regions vulnerable to stress. Based on a recent GWAS across multiple psychiatric disorder, multiple pleiotropic risk genes were identified.^34^ Furthermore, gene-disease associations from DisGeNet, a manually curated database, were used to assess risk gene enrichment for psychiatric, brain and non-brain diseases and non-disease traits. Enrichment was found for neuropsychiatric disorders (schizophrenia, bipolar disorder, and autism spectrum disorder) and other brain diseases (Parkinson’s disease). However, no gene enrichment was found for non-brain diseases (e.g. osteoporosis) and non-disease traits (e.g. height and waste-hip-ratio; **Figure 4**). Thus, differentially expressed genes in the stress network are predominantly involved in genes relevant for stress-related diseases but not in non-brain-related disorders and traits.

**Figure 4.**
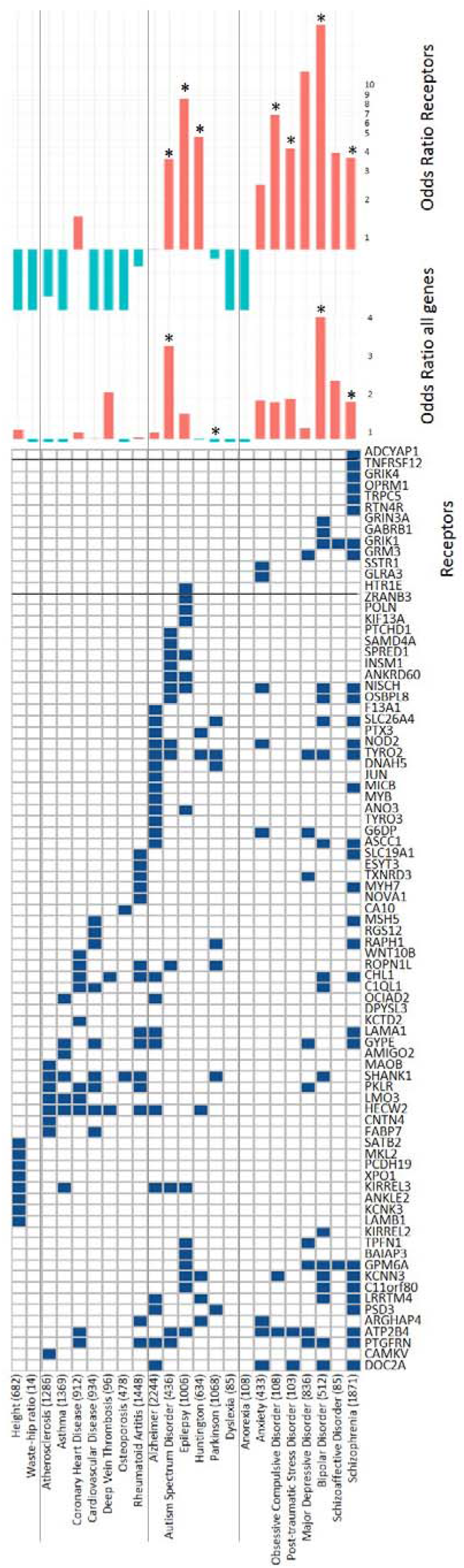
Differentially expressed genes in the stress network and stress-related psychiatric disorders. Disease risk gene enrichment was performed for the differentially expressed genes. The diseases are clustered as non-brain related disease, brain disease and psychiatric disease. As a none-disease-associated set of genes, waste-hip ratio and height were used. Numbers between the parenthesis indicate the number of genes known to be associated with the disease, based on DisGeNet. The effect size of the gene enrichment is presented at the middle part of the figure and considered significant when the BH-adjusted p-value < 0.05 (*). Blue bars mean depletion of genes, whereas red bars indicate enrichment of genes in a trait. ORs for the amount of receptors in the set of differentially expressed genes are depicted for every trait (shown on the right side of the graph).

Interestingly, the set of 201 differentially expressed genes in the stress network included a considerable number of receptors. Apart from their use as markers for specific cell types (e.g. ADCYAP1, GABRB1, SSTR1, and TNFRSF12A), these are important for signaling in the brain (e.g. ADCYAP1 modulates glutamatergic signaling^35^ and the HPA-axis response^36^) and some of them are known to be involved in the regulation of stress.^36, 37^ Therefore, we assessed whether there were more receptors in our set of genes associated to psychiatric disorders than you would expect by chance. We found higher odds ratios for brain and psychiatric disorders, with the biggest effect sizes in psychiatric disorders (**Figure 4**). Effect sizes for receptor enrichment in Epilepsy (p-value < 1*10^−5^, OR = 9.07), Huntington (p-value = 0.007, OR = 5.18), Obsessive Compulsive Disorder (p-value = 0.004, OR = 7.18), Major Depressive Disorder (p-value = 1*10^−5^, OR = 13.5), Bipolar Disorder (p-value = 1*10^−5^, OR = 26.8), and Schizophrenia (p-value = 0.03, OR = 3.82) were significant.

### 3.4 Cortisol sensitivity of the stress network

The enrichment of the neuronal GO terms in our set of genes and the association with stress-related diseases indicates that the differentially expressed genes in the stress network are relevant for stress and may be responsive to the pivotal stress hormone cortisol. To investigate glucocorticoid sensitivity, we compared our list of differentially expressed genes with genes that show a DNA binding site for the glucocorticoid- and/or mineralocorticoid receptors (GR and MR) in the rat hippocampus by Chromatin Immunoprecipitation sequencing after stimulation with the endogenous steroid corticosterone.^33^ Differentially expressed genes that showed DNA binding loci for both the GR and MR are: *SLC26A4, IGSF11, GRIK4, SCN3B, GABRB1, CCDC85A, KIRREL3, HECW2* and *PKIA* (9/459 genes with binding sites, BH-adjusted p-value for enrichment = 0.038). There was no significant enrichment for either GR binding (*OPRM1, PTER, FUT9, EPB41L4B, PKLR, GPM6A,* and *GSC;* 7/704 genes with binding sites, BH-adjusted p-value = 0.98) or MR binding exclusively (*ADCYAP1, LAMA1, CNTN4, XPO1, RGS12, MGRN1, CHST15,* and *ANKLE2*; 8/1247 genes with binding sites, BH-adjusted p-value = 0.18). These results indicate that differentially expressed genes in brain the stress network are enriched for DNA-binding loci of that (in the rat) can be bound by both the GR and the MR.

## 4. DISCUSSION

In this study, we identified genes and pathways in the cortical stress network based on an fMRI-based study involving acute stress exposure. By combining fMRI data to gene expression data, we found 201 differentially expressed genes involved in neuronal processes and enriched in stress-related psychiatric disorders. Moreover, the enriched genes included several neuropeptides and neurotransmitter receptors with regulation by both the GR and MR and substantial links to HPA-axis activity. This gene set uncovered by combining human gene expression and neuroimaging results give important new insights into the putative neural populations and mechanisms underlying stress vulnerability in humans.

Our results point to the involvement of (cortical) cell type markers in differential stress reactivity. For example, we found enrichment for some astrocyte markers, which among others modulate glutamate metabolism and transmission^38^, and there is evidence from both human and rodent models that they may play a role in stress-related disorders.^39^ Moreover, the differentially expressed genes are in general highly expressed in excitatory glutamatergic compared to inhibitory GABAergic neurons ^40^ Thus, glutamate signaling seems to be involved in a more global level, whereas GABA-related mechanisms that may underlie differential reactivity to stress are limited to a specific subset of GABA-ergic neurons. Specific targeting of these GABAergic populations, based on their receptor repertoire, may help to separate primary from secondary changes in the cortical circuitry.

For genes that do not represent specific neuronal subtypes, changed expression levels may reflect differential responsiveness based on more generic signaling pathways. This may, in particular, be the case for the identified stress-related genes with a genetic association to schizophrenia. *OPRM1* encodes for a mu-opioid receptor, which has been shown to interact with glutamate to adapt to chronic drug abuse, a stress-related disorder.^41^ Moreover, mu-opioid receptors are known to modulate the HPA-axis.^42^

Genes with high expression levels in the regions vulnerable to stress include neuropeptides and neurotransmitter receptors, which may be directly targeted to modify the activity of these brain regions. *SST1* codes for the somatostatin receptors, a neuropeptide produced in the hypothalamus. This neuropeptide is known to attenuate the stress response, by counteracting CRH signaling via the SST1 receptor.^43^ Also a number of serotonergic, GABAergic, and glutamatergic receptors are differentially represented in the stress network. All these factors may well have a role in regulating neuronal network activity during maladaptive stress responses.^44–46^ Of note, the excitatory 5-HT1E receptors are overrepresented in brain regions that failed to shut off after stress in at-risk subjects. Antagonism of 5-HT2A is common between several antipsychotic and antidepressant drugs, and normalizing the activity of these brain regions after stressor exposure may be part of their therapeutic mechanism. However, the exact function of the 5-HT1E receptors are unknown, but *HTR1E* is a candidate gene for several stress-related disorders.^47–49^

The enrichment analysis of gene ontology terms suggests that the list of differentially expressed genes play a role in stress vulnerability and risk for psychiatric disorders. For example, prenatal chronic stress has consequences on nervous system development as shown in mice.^50–52^ Moreover, disruption of neuronal plasticity^53^ is induced by a prolonged stressor and is a common symptoms of stress-related psychiatric disorders.^54^

Furthermore, we found that differentially expressed genes in the stress network are enriched for DNA-binding loci of both the GR and the MR based on experimental data in rats. GR is thought to facilitate recovery and adaptation in the aftermath of stress^55^ and polymorphisms as well as post-translational modifications alter susceptibility for stress-related psychiatric disorders.^56, 57^ The MR has been shown to facilitate stress reactivity.^58^ The link with GR and MR suggests that it related to factors related to systemic adaptations, even though we cannot know to what extent these loci actually reflect target genes.

We found a significant overlap between the genes found to be differentially expressed in the whole stress network and those found to be differentially expressed in the region-specific stress-network analysis of which some are known to be involved in stress-related phenotypes.^59, 60^ The differences between the results obtained in these experiments can be partially explained by the fact that the AHBA samples were collected using bulk sequencing which does not allow the detection of differences across individual cell populations.^61^ With the increasingly availability of single cell data we will have enough resolution to detect more subtle differences within the cortex, but for now, human brain single cell data is very limited.^24, 62–64^ Moreover, previous studies has shown that structures within the cortex are relatively similar in terms of gene expression.^14^ Therefore, the finding of 201 differentially expressed genes, point out to a true difference in the cortical stress-network and all other cortical brain regions. The non-overlapping genes from the combined analysis of cortical and non-cortical samples might be driven by anatomical differences, although it is complex to entangle the true biological signals from anatomy-driven signals.

We do not know whether the differentially expressed genes are subject to genetic regulation and whether they show differential translational responses. Furthermore, we could not infer causality, but rather association of genes with stress-sensitivity. In this regard, it will be of considerable interest to further study the genes that have been linked to psychiatric disorders, as genetic variation may, in fact, lead to abnormal expression of the genes we identified. It will also be of interest to study epigenetic regulation of the genes of interest and gene-environment interactions.^65–68^

Given that we assessed gene expression levels in the healthy brain, it is challenging to interpret the differences in high and low expression levels and the meaning in diseased brains. High expression levels of the genes in the stress network do not necessarily mean that stress sensitivity is a result of the high gene expression *per se*. It might be the ability to regulate neurobiological processes via direct neurotransmitter and receptor signaling or the ability to indirectly regulate changes in gene expression.^69^ Moreover, we have to take into consideration that we identified genes that already show low baseline expression levels in the brain.

Another limitation of this study is that the low number of samples in some brain regions did not allow the analysis of differential expression within these regions. For example, the precuneus and the angular gyrus were underrepresented in the AHBA (n = 7 for both regions), but harbored great changes according to the fMRI signal. However, there were sufficient brain samples available from the AHBA to analyze brain regions vulnerable to stress altogether. Moreover, the stress-network we defined was based on 78 males. Given the relatively small sample size, replication in a bigger independent cohort should be awaited. Furthermore, the six donors were five males and one female. It is important to take donor’s sex effect into account, since there is a sex difference in the development and symptoms of stress-related diseases^70, 71^. Therefore, we checked whether gene expression levels were different for the female donor compared to the male donors. We did not find gender effects of gene expression levels of the differentially expressed genes. To maximize the number of samples, we decided to include the female donor in our analyses. It has to be taken into account, however, that the outcome of the performed task might be different across the genders.^72^ This implies that our results cannot be generalized over the whole population, but are rather reflective for males, since the stress-network we identified could also be male-specific. We also acknowledge that we do not know how the stress network would look like in individuals at risk for other psychiatric disorder. Moreover, brain regions differentially activated by acute stress are specific for the emotion processing task. Therefore, we might have missed some relevant brain structures, and thus genes, that might have become active during another task under stressful conditions. Lastly, the stress network that was used in this paper was based on data from siblings of patients with schizophrenia. Even though stress is a transdiagnostic factor and relevant for all psychiatric disorders^73^, we cannot directly extrapolate the stress network to other psychiatric disorders such as depression and bipolar disorder. There is increasing research into risk groups for these disorders, but to our knowledge, direct comparisons on brain-related stress sensitivity between (risk groups of) across psychiatric disorders are lacking.

To our knowledge, this is the first study to map gene expression atlases to task-based fMRI data in order to identify the molecular mechanisms underlying human stress reactivity in relation to the risk to develop psychiatric disorders. Here, we show that this method can aid in disentangling the molecular underpinnings of specific tasks and traits. We showed that genes possibly underlying stress reactivity are also associated with neuronal cell type markers (e.g. glutaminergic excitatory neurons), stress-related disease, GR and MR responsiveness and HPA-axis activity. We identified several neuropeptides and receptors as important players. These identified systems are not only important to understand the underlying mechanisms of stress vulnerability, but can also be used to develop new drug targets. Therefore, identification of novel drug targets involved in stress vulnerability would be of great interest for the development of new therapies in stress-related psychopathology.

## FUNDING AND DISCLOSURE

The authors declare no competing financial interests and have no things to disclose.

## ACKNOWLEDGEMENTS

The authors would like to thank Sjoerd Huisman for his advice and help on some of the statistical aspects of the study and Martijn van den Heuvel for his advice on the manuscript.

## Notes

### Competing Interest Statement

The authors have declared no competing interest.

