## Supplementary figures and images for "Molecular characterization of the stress network in the human brain"

### Figure S1

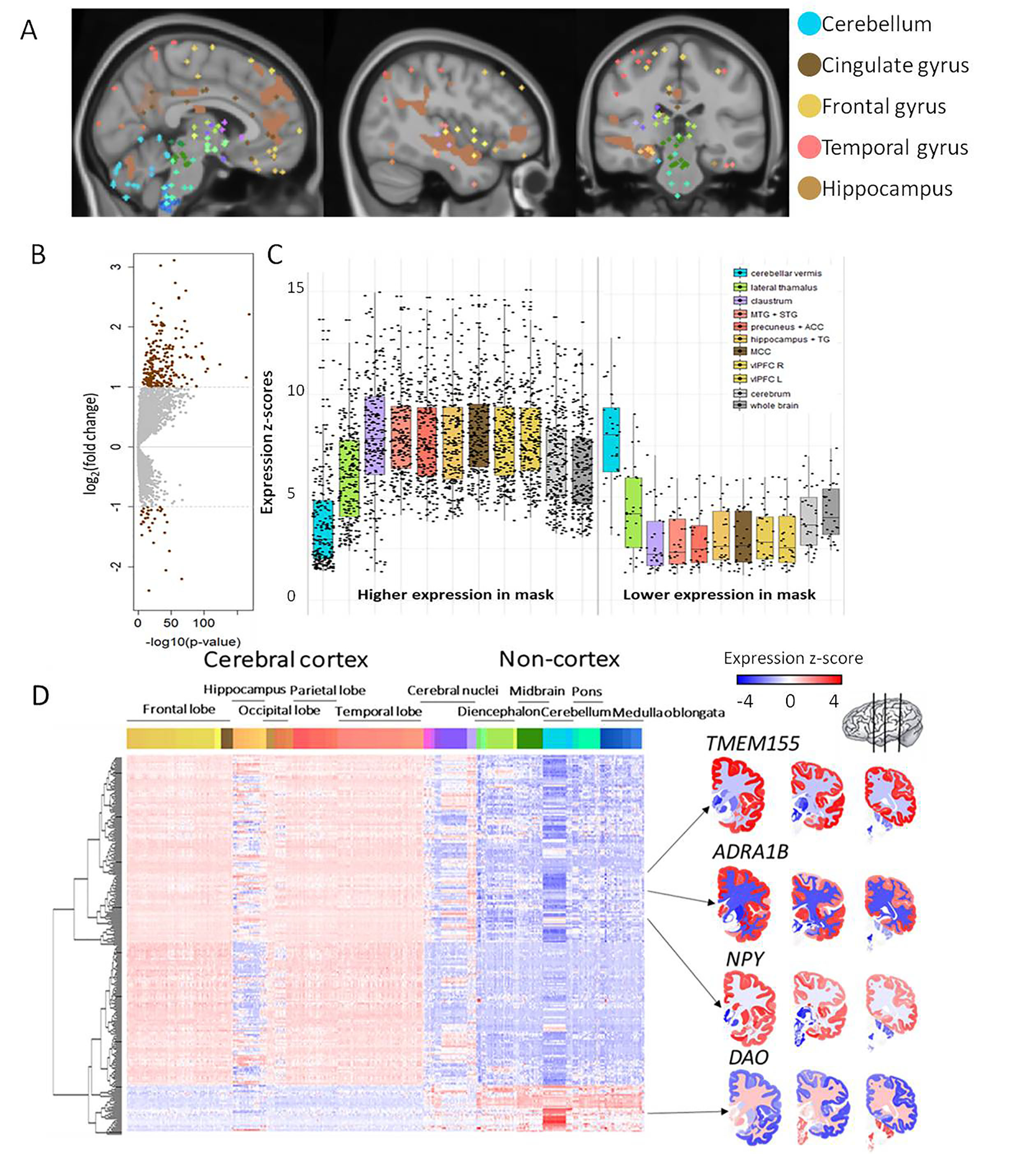

### Figure S2

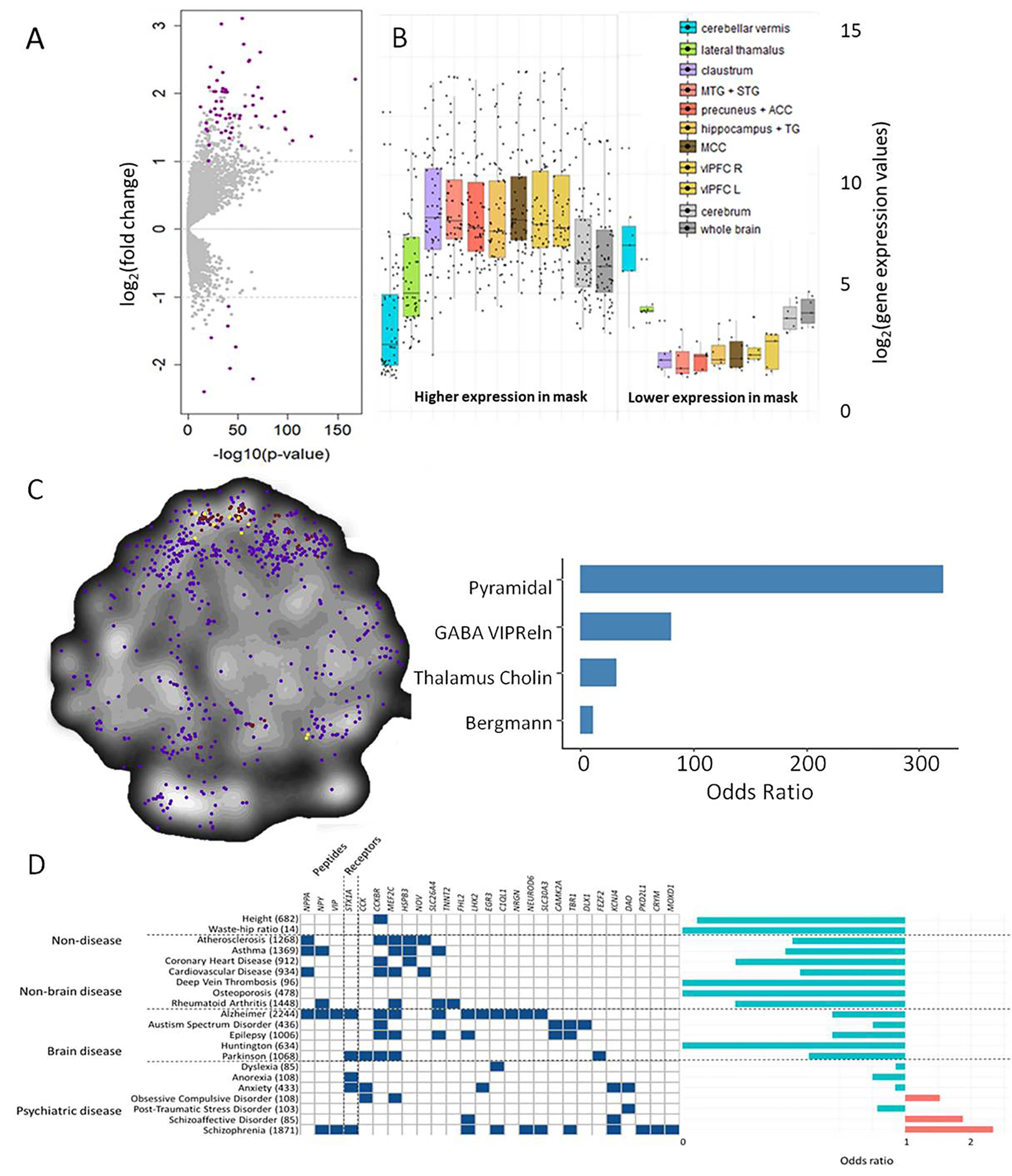

### Figure S3

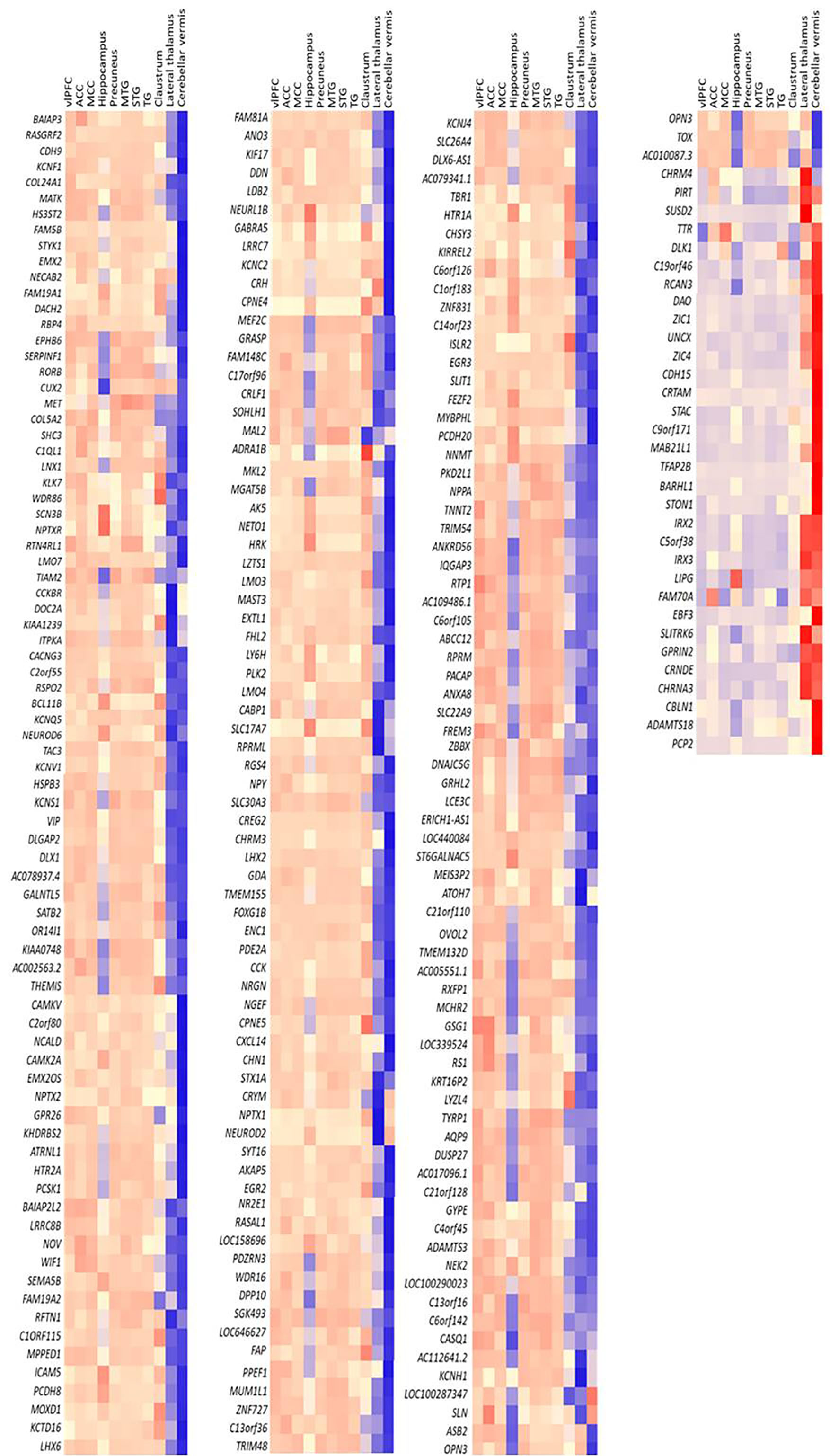

### Figure S4

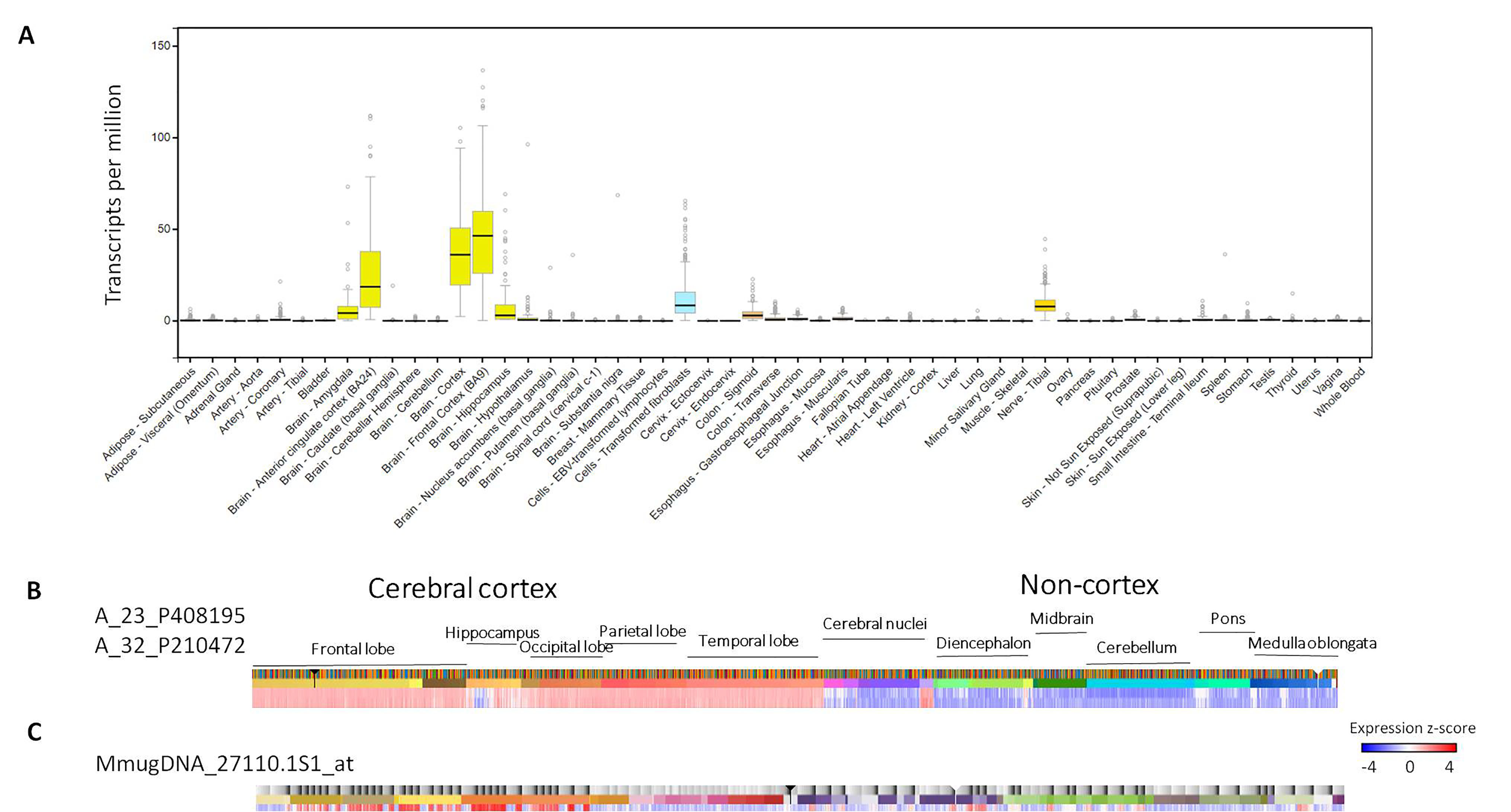

### Figure S5

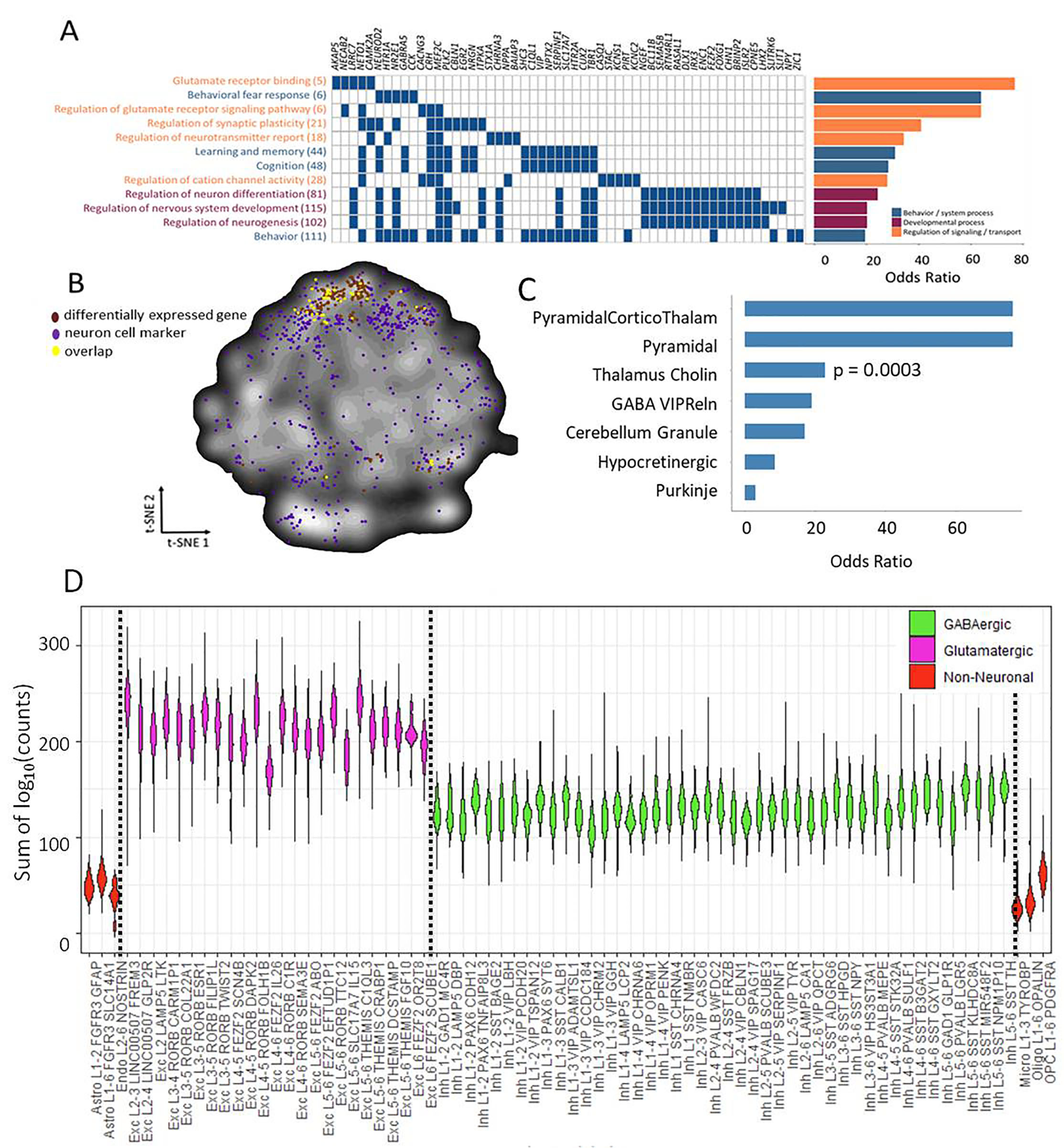

### Figure S6

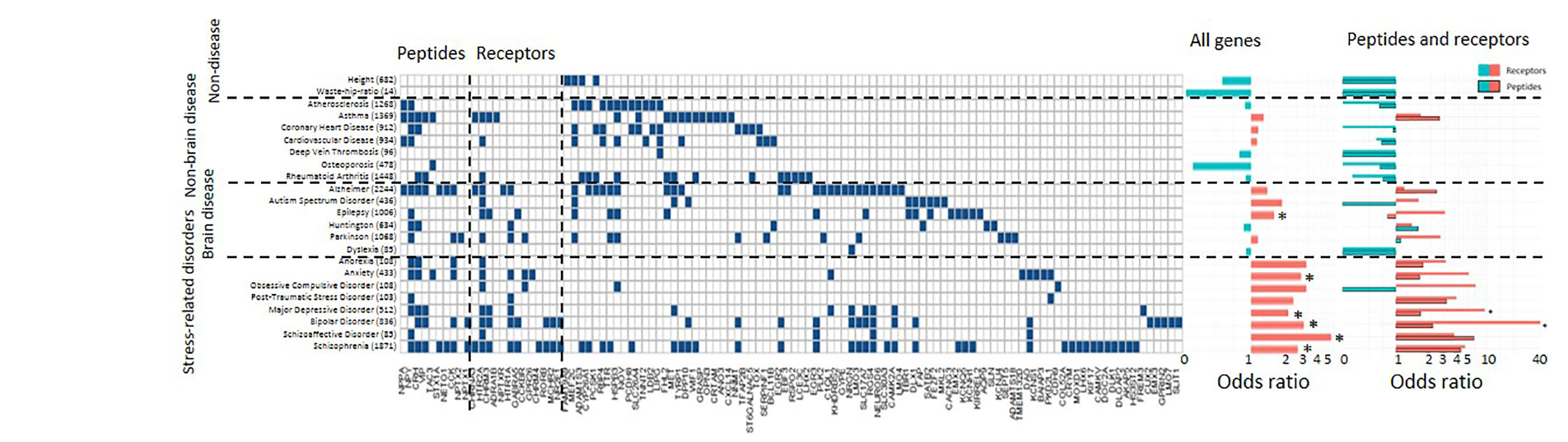

### Figure S7

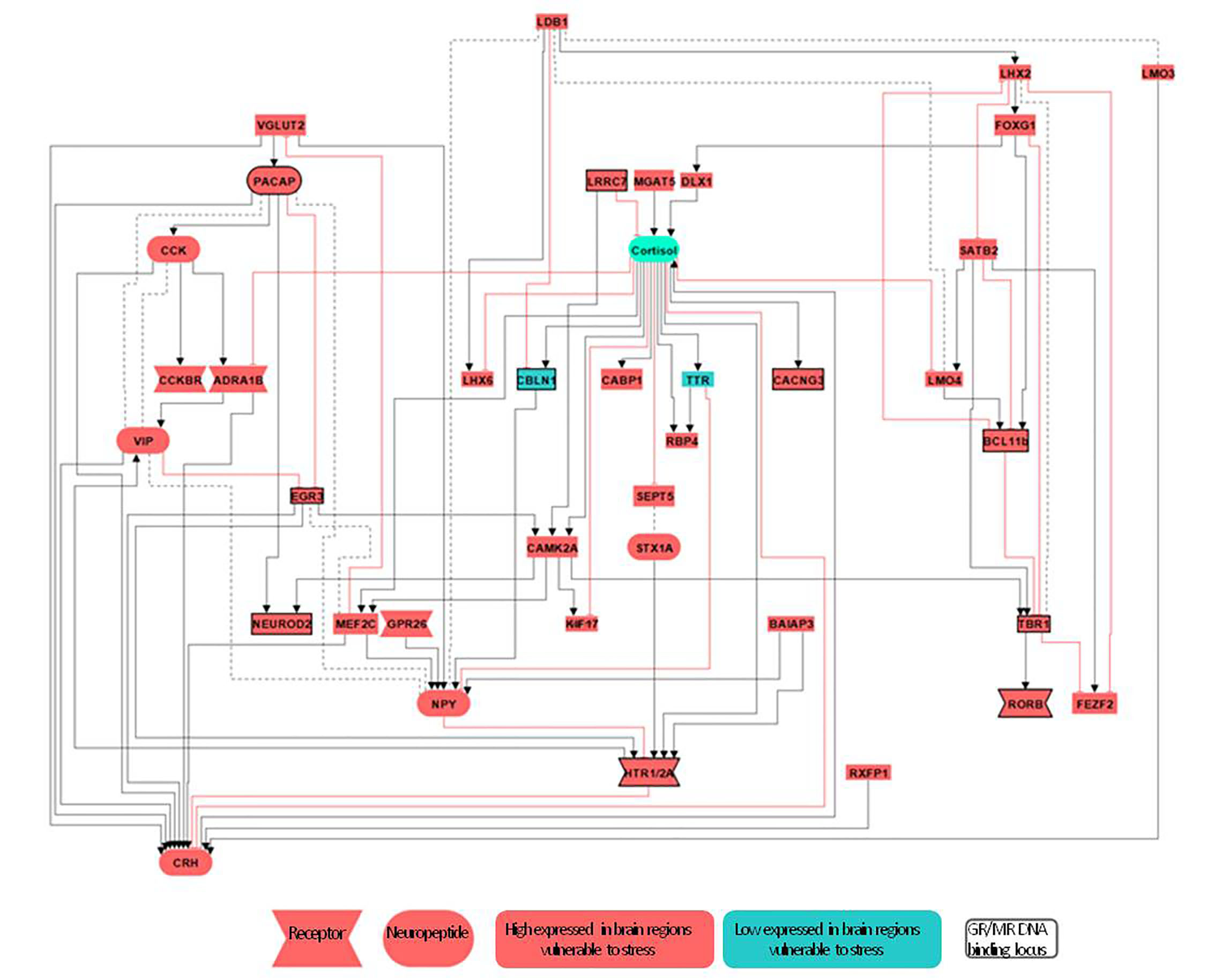
