## supplementary text for "Molecular characterization of the stress network in the human brain"

**Differentially expressed genes in the stress network**

We identified the gene expression signatures of the stress network with altered stress-induced activity by determining which genes are differentially expressed in the stress network compared to the rest of the brain minus the cerebellum. Using a meta-analysis approach to combine results across all donors of the AHBA (n=6), we identified 261 differentially expressed genes (BH-adjusted p < 0.05 and log_2_(fold-change) > |1|, **Figure S1; Table S2**). Among those genes, 229 were higher expressed, while the other 32 genes were lower expressed in the stress network compared to the rest of the brain minus the cerebellum. Using a bootstrapping approach (see Methods, Main Text), we found the identified set of genes to be highly robust to the imbalance between the number of AHBA samples inside and outside the stress network (61% of our initial 261 genes were differentially expressed in at least 95% of all 1000 iterations).

As a sensitivity analysis, we next selected genes that were differentially expressed (BH-adjusted p < 0.05 and log_2_(fold-change) > |1|) in at least five out of the six donors. This more stringent approach resulted in a smaller set of 63 differentially expressed genes (**Figure S2**). The smaller set of 63 genes was entirely contained within the larger set of 261 genes obtained by the meta-analysis, therefore we choose to perform subsequent analyses on the larger gene set.

The differentially expressed genes in the brain regions vulnerable to stress showed an opposite expression pattern in the cerebellum compared to the rest of the brain (**Figure S1B**), supporting our approach of excluding the cerebellum from our analysis (see Methods).

To assess whether our results are driven by the higher representation of cerebral cortex samples in the brain regions vulnerable to stress (109 out of 127; 91%) compared to the rest of the brain (1,950 out of 3,225; 60%), we identified differentially expressed genes within each anatomically-defined region (hippocampal formation, cerebellum, frontal gyrus, temporal gyrus and cingulate gyrus) separately (**Table S3**). Due to the small sample size, it was not possible to assess differential gene expression between brain regions vulnerable to stress compared to the rest of the brain minus the cerebellum within each anatomically-defined region (**Table S4**).

The differentially expressed genes consistently showed high expression values in the cortex but not the hippocampus, and mostly low expression levels in non-cortical areas (**Figure S1D and Figure S3**). The two most differentially expressed genes in the stress-specific brain regions are D-amino acid oxidase (*DAO)* (BH-adjusted p-value = 4.99*10^-17^, log_2_(FC) = -2.40) and Transmembrane protein 155 (*TMEM155)* (BH-adjusted p-value = 4.90*10^-55^, log_2_(FC) = 3.11)*. TMEM155* is highly expressed in the brain, specifically in the cerebral cortex and claustrum (**Figure S4**).^35^ Its biological function, however, remains unknown, largely because there are no orthologs in the Mus musculus or the Rattus norvegicus. *TMEM155* orthologs are, however, found in chimpanzee^36^ and rhesus macaque^37^, with similar brain expression patterns between the human and macaque (**Figure S4B and S4C**).

**Functionality and cell-type specificity of differentially expressed genes in the stress network**

A GO term enrichment analysis was performed to assess whether the differentially expressed genes in the stress network are enriched for specific functions. The differentially expressed genes were enriched for GO terms involved in neuronal development and neurogenesis, synaptic signal transmission, learning and memory, behavior and glutamate receptor signaling (**Figure S5A and Table S2**). Genes involved in most processes based on GO terms (at least assigned to five out of twelve GO terms) include *CRH, MEF2C, NETO1, PLK2, NEUROD2, NR2E1, SERPINF1, CHRNA3, ISLR2, CUX1* and *TBR1*. Enrichment analysis for cellular components indicated that the proteins coded by the differentially expressed genes were mainly found at the synapse, reflecting the high expression of the genes in the synapse-dense cerebral cortex. To control for the case that the results of the GO term enrichment analysis are solely driven by genes already known to be associated with stress-related disorders, the analysis was also performed after exclusion of 58 genes associated to stress related disorders. We found still very similar results, indicating that differentially expressed genes in the brain regions vulnerable to stress are predominantly involved in genes relevant for stress-related diseases but not in non-brain-related disorders and traits.

Next, we identified the specific cell types underlying the differential gene expression levels in the stress network using enrichment analysis of cortical cell-type markers.^23^ Enrichment was found for neuronal cell markers (p = 0.0414, BH corrected), including *WIF1, CHRNA3, TAC3, VIP, C1QL1, SLITRK6, NEUROD6, KCTD16, TBR1, CREG2* and *DLX1*. The list of differentially expressed genes included a few astrocytes markers (*AQP9, EMX2, KIRREL2, NR2E1* and *LHX2*) and oligodendrocytes (*EBF3*), but was not significantly enriched (p = 0.4202; p = 0.5267, BH corrected). Moreover, we found that neuronal markers showed a partially overlapping distribution in a t-Distributed Stochastic Neighbor Embedding (t-SNE) map of all genes across the whole brain as the differentially expressed genes in, indicating that neuronal markers and the differentially expressed genes show the same expression patterns across cortical areas (**Figure S5B**) and thus differential activity may depend on neuronal gene expression.

We further assessed whether the set of differentially expressed genes was particularly expressed in a neuronal subtype. Interestingly, five out of the 22 known thalamic cholinergic neuron markers were found in the set of the differentially expressed genes (*CHRNA3* (BH-adjusted p-value = 6.6*10^-6^, log_2_(FC) = -1.47)*, IRX2* (BH-adjusted p-value = 1.31*10^-^42, log2(FC) = -2.03)*, NPPA* (BH-adjusted p-value = 1.85*10^-34^, log2(FC) = 1.89)*, TYRP1* (BH-adjusted p-value = 3.3*10^-14^, log_2_(FC) = 1.1) and *ZIC4* (BH-adjusted p-value = 7..57*10^-8^; log_2_(FC) = -1.24), BH-adjusted p-value = 3.3*10^-4^; **Figure S5C**). This finding in thalamic cholinergic neuron markers is independent of the number of thalamic samples inside and outside the brain regions vulnerable to stress, showing that this result is not biased by the number of thalamic samples involved in the maladaptive stress response. Additionally, some of the differentially expressed genes are reported to be part of the molecular fingerprint of particular cell types in the human cortex.^38^ For example, *RORB* represents a subclass of excitatory neurons present in the cortex and was modestly enriched in differentially activated brain regions. *VIP* represents one of the four main subclasses of inhibitory cortical neurons and was quite substantially overrepresented (BH-adjusted p-value = 5.73*10^-69^, log_2_(FC) = 1.66). Markers for smaller subsets of inhibitory neurons included overrepresentation of *NPY* (a small set of deep projecting inhibitory layer 3 neurons; BH-adjusted p-value = 3.53*10^-25^, log_2_(FC) = 1.60) and *SERPINF1* (a subset of inhibitory neurons in layer 1; BH-adjusted p-value = 7.83*10^-37^, log_2_(FC) = 1.27), and underrepresentation of *CBLN1* (a subset of VIP expressing inhibitory neurons in layer 4; BH-adjusted p-value = 1.85*10^-11^, log_2_(FC) = 1.20). Using a human-specific single cell RNA-sequencing data of the medio-temporal gyrus^25^, we found the differentially expressed genes to be mainly enriched in glutamatergic excitatory neurons compared to GABAergic and non-neuronal cells, using a Wilcoxon rank test (p-value = 2.2*10^-^16, **Figure S5D**).

**Differentially expressed genes in stress network are associated to stress-related diseases**

We hypothesized that the differentially expressed genes in the stress network would be associated to the genetic background of psychiatric disorders, particularly for stress-related brain disorders, as stress plays a major role in the development of these disorders. Using genetic variants from GWAS of the Genomics of Psychiatry Consortium^26, 27^, we assessed whether schizophrenia-associated risk loci are enriched in the set of differentially expressed genes. Indeed, schizophrenia risk genes were enriched in the differentially expressed genes in the stress network (Fisher Exact test, BH-adjusted p-value = 2.4*10^-3^). The schizophrenia risk genes *KCNV1, DOC2A, NGEF, NRGN, MEF2C, BCL11b, SATB2, FAM5B, CHRM3, CHRM4* and *TAC3* were present in our differentially expressed genes, and all except one (*CRHM4*) were higher expressed in the brain regions vulnerable to stress. Based on a recent GWAS across multiple psychiatric disorder, multiple pleiotropic risk genes were identified.^39^ Differentially expressed genes in the stress network were significantly enriched in the set of pleiotropic risk genes for psychiatric disorders (Fisher Exact test, BH-adjusted p-value = 0.0065, OR = 30.3), including *KCNQ5, BCL11b, NGEF, CHRNA3, NRGN,* and *CHRM4*. Furthermore, gene-disease associations from DisGeNet, a manually curated database, were used to assess risk gene enrichment for psychiatric, brain and non-brain diseases and non-disease traits. Enrichment was found for neuropsychiatric disorders (schizophrenia, schizoaffective disorder, major depressive disorder, bipolar disorder, and anxiety disorders) and other brain diseases (epilepsy). However, no gene enrichment was found for non-brain diseases (e.g. osteoporosis) and non-disease traits (e.g. height and waste-hip-ratio; **Figure S6**). Thus, differentially expressed genes in the stress network are predominantly involved in genes relevant for stress-related diseases but not in non-brain-related disorders and traits.

Interestingly, the set of 261 differentially expressed genes in the stress network included a considerable number of neuropeptides and receptors. Apart from their use as markers for specific cell types (e.g. NPY, VIP, and CCK^38^), these are important for signaling in the brain (e.g. NPY stimulates the release of CRH in the hypothalamus^40^) and some of them are known to be involved in the regulation of stress.^41, 42^ Therefore, we assessed whether there were more neuropeptides and receptors in our set of genes than you would expect by chance. For both the neuropeptides and the receptors, we found higher odds ratios for brain and psychiatric disorders, with the biggest effect sizes in psychiatric disorders (**Figure S6**). Effect sizes for receptor enrichment in major depressive disorder and bipolar disorder were significant (p-value = 0.0003, OR = 1.86 and p-value < 0.00001, OR = 2.57, respectively).

**Cortisol sensitivity of the stress network**

The enrichment of the neuronal GO terms in our set of genes and the association with stress-related diseases indicates that the differentially expressed genes in the stress network are relevant for stress and may be responsive to the pivotal stress hormone cortisol. To investigate glucocorticoid sensitivity, we compared our list of differentially expressed genes with genes that show a DNA binding site for the glucocorticoid- and/or mineralocorticoid receptors (GR and MR) in the rat hippocampus by Chromatin Immunoprecipitation sequencing after stimulation with the endogenous steroid corticosterone.^34^ Differentially expressed genes that showed DNA binding loci for the GR exclusively are: *STON1, HS3ST2, HTR2A, ZNF831, CACNG3, NPTX2, EPHB6, LRRC7, KCNC2, HTR1A, NETO1, CYP26A1, NCALD, EMX2, CXCL14* and *RORB* (16/704 genes with binding sites, BH-adjusted p- value = 2.97*10^-2^). The differentially expressed genes *HSPB3, EGR3, NEUROD2, TBR1, SLC26A6, SLIT1, PCDH20, MAST3, BCL11b, SCN3B, TFAP2B, MAB21L1* and *CHN1* have DNA binding loci for both the GR and the MR (13/459 genes with binding sites, BH-adjusted p-value = 9.8*10^-3^). There was no significant MR DNA binding loci enrichment (*IRX2, IRX3, PIRT, CBLN1, KCNV1, KCNS1, ICAM5, FHL2, DLGAP, ST6GALNAC5, RFTN1, KCNQ5, KCNH1* and *LMO7*; 14/1247 genes with binding sites, BH-adjusted p-value = 0.46). These results indicate that differentially expressed genes in brain the stress network are enriched for DNA-binding loci of GR but not MR, further consolidating the relevance of these brain regions in aftermath of the responses to acute stress when the GR plays a dominant role.

**Putative molecular pathway as the basis for inter-individual differences in stress reactivity**

To create a more comprehensive possible pathway for the molecular mechanisms underlying human stress reactivity, we performed a thorough literature search on PubMed for all differentially expressed genes to assess whether previous studies have found these genes to be regulated by or regulating the HPA-axis. We found that 53 of our differentially expressed genes were earlier described to interact with the HPA-axis (**Table S5**), with a subset of 36 genes reported to interact with each other (F**igure S7, Table S5**). A putative pathway with factors involved in stress reactivity was build based on these 36 genes (**Figure S7**).

**Table S3 Differentially expressed genes in the cerebellum and hippocampus**

|  | Gene | BH-adjusted p-value | Log_2_(FC) |
| --- | --- | --- | --- |
| **Cerebellum** | *SLN* | 1.7*10^-3^ | 1.13 |
| (5 samples inside and 78 samples outside | *CCDC155* | 1.3*10^-3^ | 1.01 |
| the stress-vulnerability related brain | *SEMA3C* | 4.0*10^-3^ | 1.01 |
| regions) | *NTNG1* | 1.7*10^-3^ | -1.18 |
|  | *CHRNB3* | 2.1*10^-3^ | -1.35 |
|  | *SHB* | 6.0*10^-4^ | -1.40 |
| **Hippocampus** | *ZMAT4* | 1.04*10^-10^ | 1.30 |
| (19 samples inside and 26 samples outside | *SLC17A6* | 1.6*10^-4^ | 1.11 |
| the stress-vulnerability related brain | *PACAP* | 2.35*10^-5^ | 1.05 |
| regions) | *IGSF3* | 1.01*10^-7^ | 1.03 |
|  | *DMKN* | 5.07*10^-9^ | -1.12 |
|  | *IQGAP3* | 4.3*10^-3^ | -1.15 |
|  | *SCGN* | 5.57*10^-9^ | -1.48 |

|  | Donor 1 | Donor 2 | Donor 3 | Donor 4 | Donor 5 | Donor 6 |  | **Not in mask** | Note |
| --- | --- | --- | --- | --- | --- | --- | --- | --- | --- |
| Superior frontal gyrus | 4 | 6 | 1 | 1 | 3 | 3 |  | 3 | Cortex |
| Middle frontal gyrus | 1 | 1 | 1 | 0 | 1 | 1 |  | 5 | Cortex |
| Inferior frontal gyrus | 2 | 3 | 0 | 1 | 0 | 0 |  | 7 | Cortex |
| Parolfactory gyrus | 0 | 1 | 0 | 0 | 0 | 0 |  | 1 | Cortex |
| Superior rostral gyrus | 1 | 1 | 0 | 0 | 0 | 0 |  | 1 | Cortex |
| Supramarginal gyrus | 1 | 1 | 0 | 1 | 2 | 0 |  | 4 | Cortex |
| Angular gyrus | 0 | 0 | 1 | 0 | 1 | 1 |  | 4 | Cortex |
| Precuneus | 1 | 0 | 1 | 0 | 0 | 1 |  | 5 | Cortex |
| Superior temporal gyrus | 4 | 2 | 1 | 1 | 0 | 2 |  | 5 | Cortex |
| Middle temporal gyrus | 7 | 1 | 1 | 0 | 0 | 0 |  | 4 | Cortex |
| Heschl’s gyrus | 0 | 0 | 0 | 1 | 0 | 0 |  | 1 | Cortex |
| Transverse gyrus | 0 | 0 | 0 | 0 | 1 | 0 |  | 1 | Cortex |
| Planum temporale | 2 | 0 | 0 | 1 | 0 | 0 |  | 1 | Cortex |
| Planum polare | 1 | 1 | 2 | 2 | 1 | 1 |  | 1 | Cortex |
| Cuneus | 2 | 0 | 0 | 0 | 0 | 0 |  | 5 | Cortex |
| Anterior cingulate gyrus | 0 | 3 | 1 | 0 | 4 | 2 |  | 2 | Cortex |
| Parietal cingulate gyrus | 0 | 0 | 1 | 2 | 1 | 1 |  | 4 | Cortex |
| Hippocampal formation | 5 | 3 | 2 | 4 | 2 | 3 |  | 26 |  |
| Claustrum | 0 | 1 | 1 | 0 | 2 | 0 |  | 0 |  |
| Caudate nucleus | 0 | 1 | 0 | 0 | 1 | 0 |  | 89 |  |
| Lateral group of nuclei | 0 | 3 | 0 | 1 | 1 | 0 |  | 0 |  |
| Cerebellum | 3 | 2 | 0 | 0 | 0 | 0 |  | 78 | Excluded |

**Table S4 Distribution of samples of the Allen Human Brain Atlas**

Red = <2 samples in the donors
Orange = <2 samples outside the mask
Green = comparison was possible, but effect sizes were too small

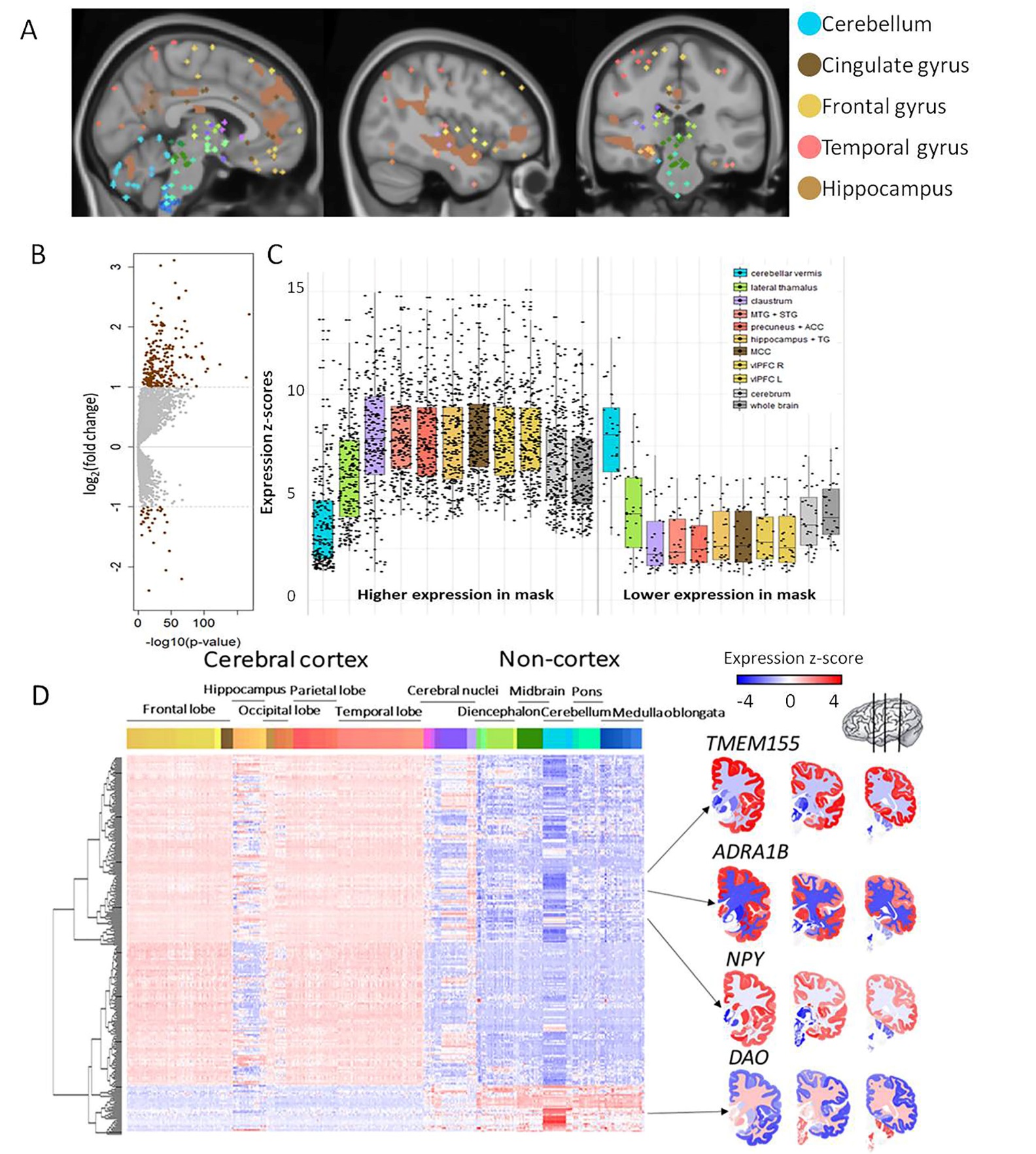
**Figure S1 Differentially expressed genes in brain regions vulnerable to stress can be identified using gene expression atlases.** (A) Brain regions in the stress network are present throughout the brain (including cerebellum, cingulate gyrus, frontal gyrus, temporal gyrus, and hippocampal formation). For the analysis, all regions were combined. (B) Differential gene expression was determined for the stress network compared to the rest of the brain minus cerebellum. Significant genes (BH-adjusted p-value < 0.05 & log_2_(fold-change) > |1|) have higher expression in the stress network. Grey dots represent non-significant genes and brown dots represent significant genes based on meta-analysis across all six AHBA donors. (C) The box plots show the expression of the higher (left) and lower (right) expressed genes compared to the rest of the brain minus cerebellum in the brain regions of interest from the stress network. (D) In the whole brain, differentially expressed genes show mostly high expression levels in the cortex and low expression levels in non-cortical brain regions. In the heatmap, each row represents a gene and each column represents a sample and all samples of the AHBA are illustrated here. On the right, coronal brain sections for the genes *TMEM155*, *ADRA1B*, *NPY* and *DAO* are presented. Colors indicates high (red) and low (blue) expression levels.

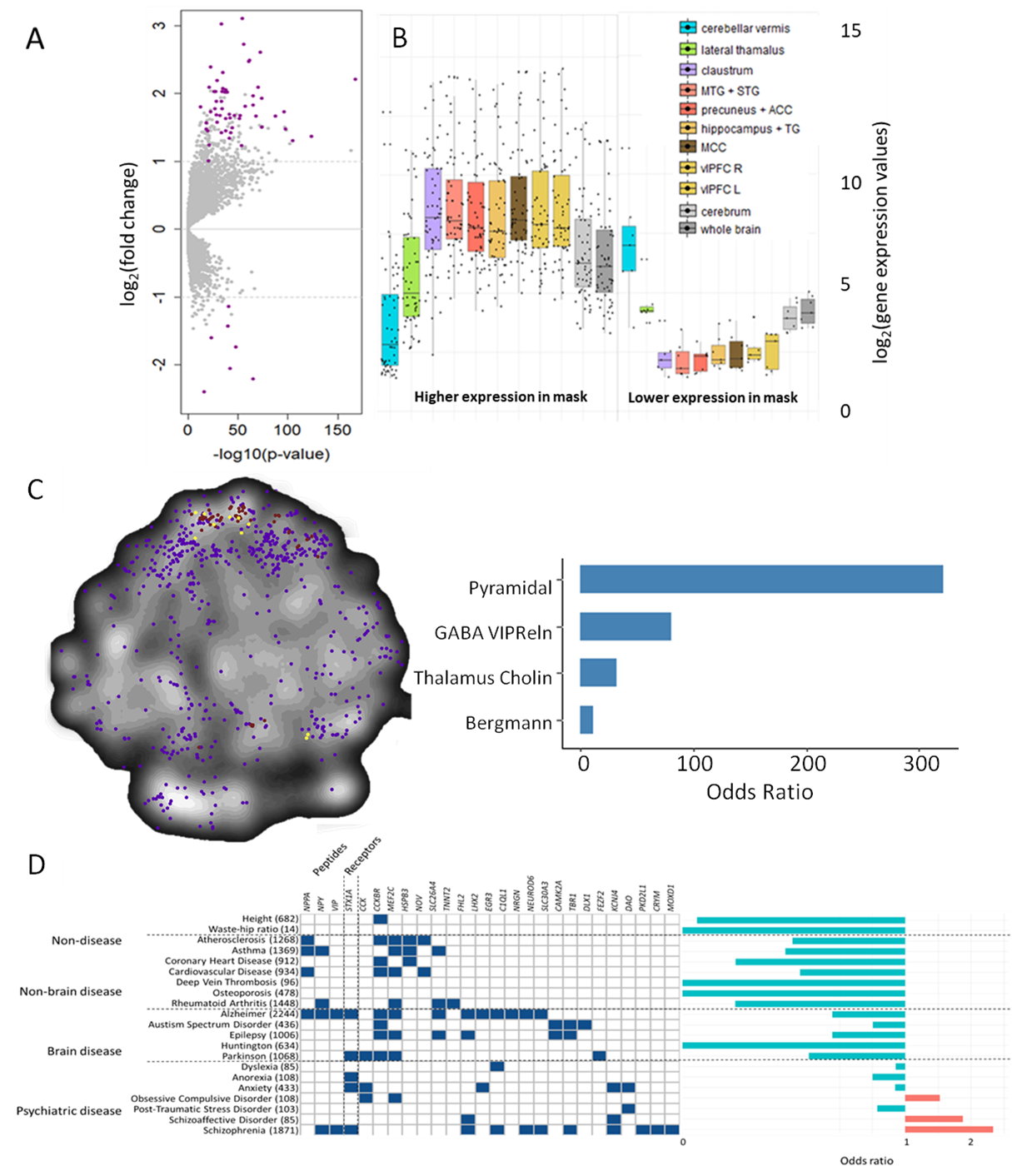

**Figure S2 Using a stricter method of selecting differentially expressed genes results in similar outcome as using a meta-analysis.** (A) Differential gene expression was determined for the stress network compared to the rest of the cerebrum for each of the six donors separately. Genes showing significance (BH-adjusted p-value < 0.05 & log_2_(fold-change) > |1|) in at least 5 out of the 6 donors were selected and are shown in purple in the volcano plot of the meta-analysis. Grey dots represent non-significant genes. The box plots on the right show the expression of the individual differentially expressed genes relative to the average gene expression in individual brain regions. (B) Differentially expressed genes (brown), neuronal marker genes (purple) and overlapping genes (pink) are plotted in a t-Distributed Stochastic Neighbor Embedding (t-SNE) plot acquired from BrainScope.nl^21^, visualized in a heatmap of genes, where close points represent genes with similar gene expression profiles. ORs for different neuronal subtypes are presented. Neuronal subtypes without a marker represented in the differentially expressed genes are not illustrated in the graph. No significant enrichment for neuronal markers was observed. (C) Disease-risk gene enrichment was performed for the differentially expressed genes. The diseases are clustered as non-brain related disease, brain disease and psychiatric disorders. As a none-disease-associated set of genes, waste-hip ratio and height were used. The numbers between the parenthesis indicate the amount of genes known to be associated with the disease based on DisGeNet. The effect size of the gene enrichment is presented on the right side of the figure, though none of them was significant.

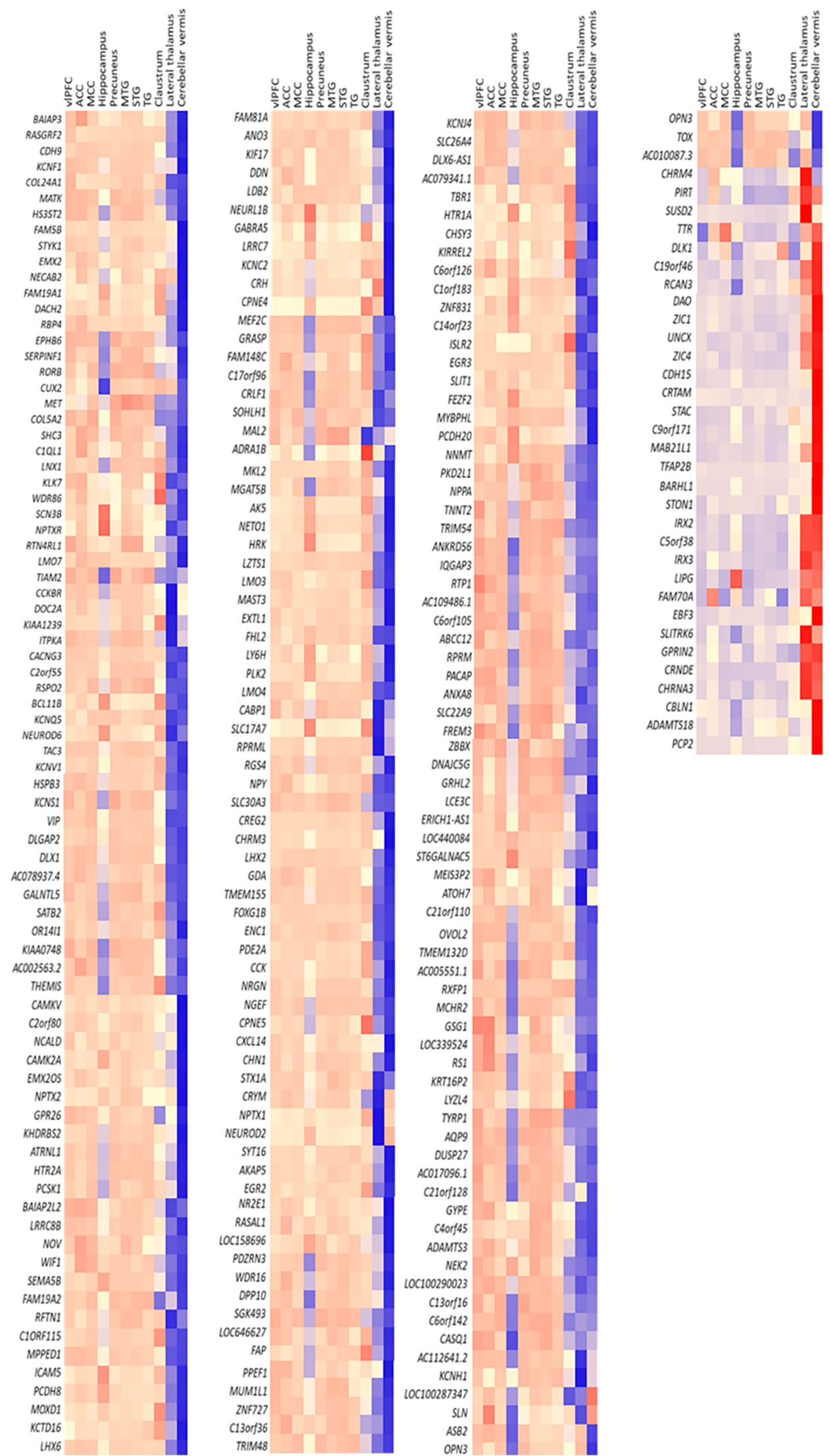

**Figure S3 Differential gene expression in the whole brain stress network.** For each brain region of the defined stress network, relative gene expression levels are shown. Colors indicates high (red) or low (blue) expression.

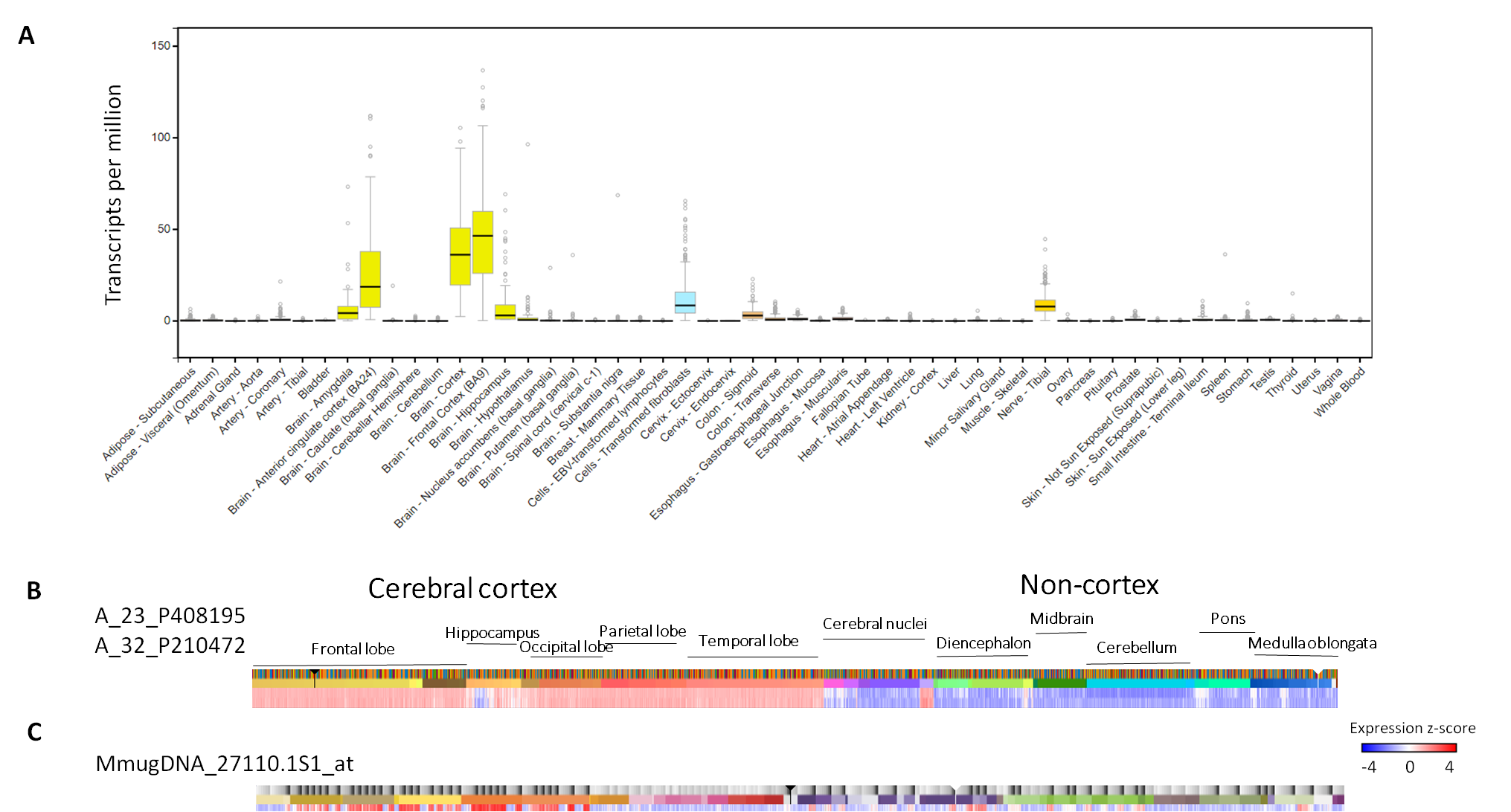

**Figure S4 *TMEM155* is highly expressed in the claustrum.** (A) In the human body, *TMEM155* has only been identified in the brain. (B) In the human brain, *TMEM155* (measured using two probes: A_23_P408195 and A_32_P210472) is highly expressed in the cortex and claustrum. (C) The expression pattern of *TMEM155* (measured using one probe: MmugDNA_27110.1S1_at) in the macaque brain resembles the human expression pattern.

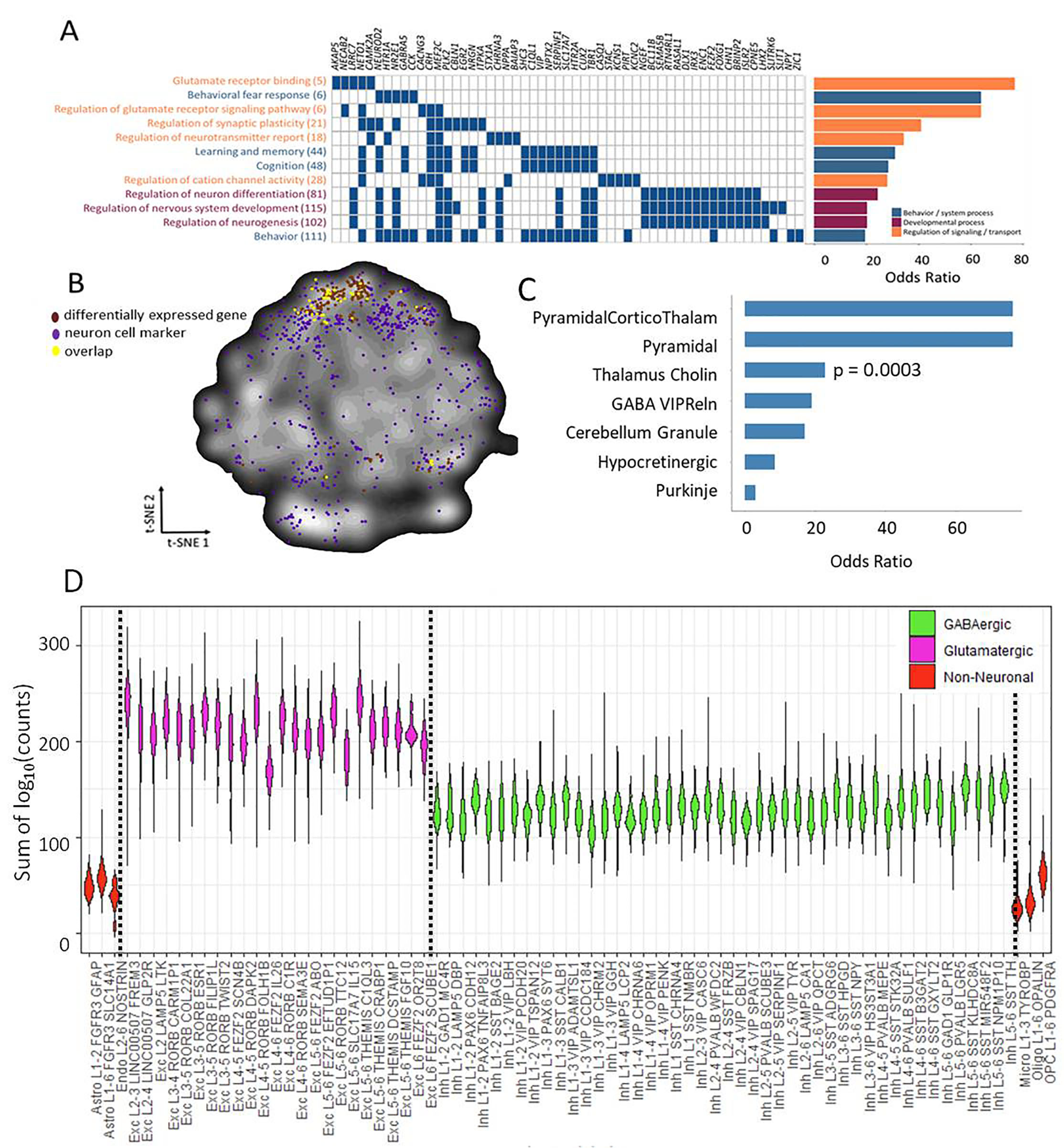
**Figure S5 Functionality of differentially expressed genes in the stress network.** (A) Differentially expressed genes annotated to one of the GO terms were assigned to multiple GO terms and thus involved in multiple processes. Between parenthesis, the total number of genes assigned to the GO term is depicted. On the right side of the graph, ORs are displayed per GO term. (B) Differentially expressed genes (brown), neuronal marker genes (purple) and overlapping genes (yellow) are plotted in a t-SNE plot generated using BrainScope.nl^21^, where points close together represent genes with similar gene expression profiles. The differentially expressed genes show a similar profile in the t-SNE plot as neuronal cell markers (purple). (C) ORs for different neuronal subtypes. ORs were considered significant when the BH-adjusted p-value < 0.05 (*). Neuronal subtypes without a marker represented in the differentially expressed genes are not illustrated in the graph. (D) The sum of the log_10_ values of the counts per gene is plotted for each cell cluster. Green clusters belong to GABAergic cells, purple clusters to glutamatergic cells and red clusters to non-neuronal cells.

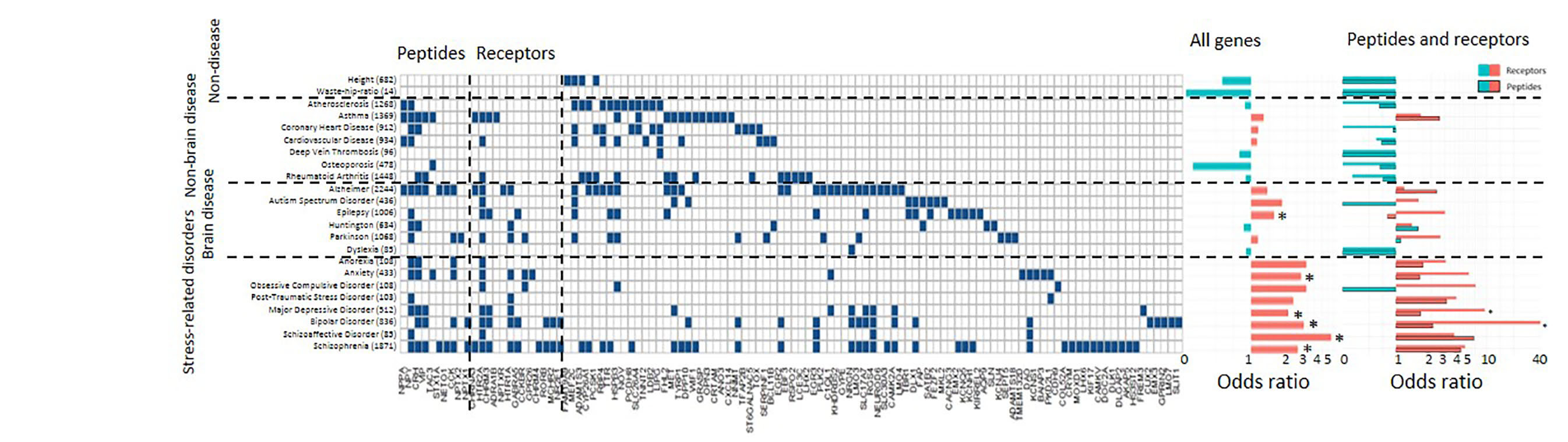
**Figure S6 Differentially expressed genes in the stress network and stress-related psychiatric disorders** Disease risk gene enrichment was performed for the differentially expressed genes. The diseases are clustered as non-brain related disease, brain disease and psychiatric disease. As a none-disease-associated set of genes, waste-hip ratio and height were used. Numbers between the parenthesis indicate the number of genes known to be associated with the disease, based on DisGeNet. The effect size of the gene enrichment is presented at the middle part of the figure and considered significant when the BH-adjusted p-value < 0.05 (*). Blue bars mean depletion of genes, whereas red bars indicate enrichment of genes in a trait. ORs for the amount of receptors and peptides (denoted by the black borders around the bars) in the set of differentially expressed genes are depicted for every trait (shown on the right side of the graph).

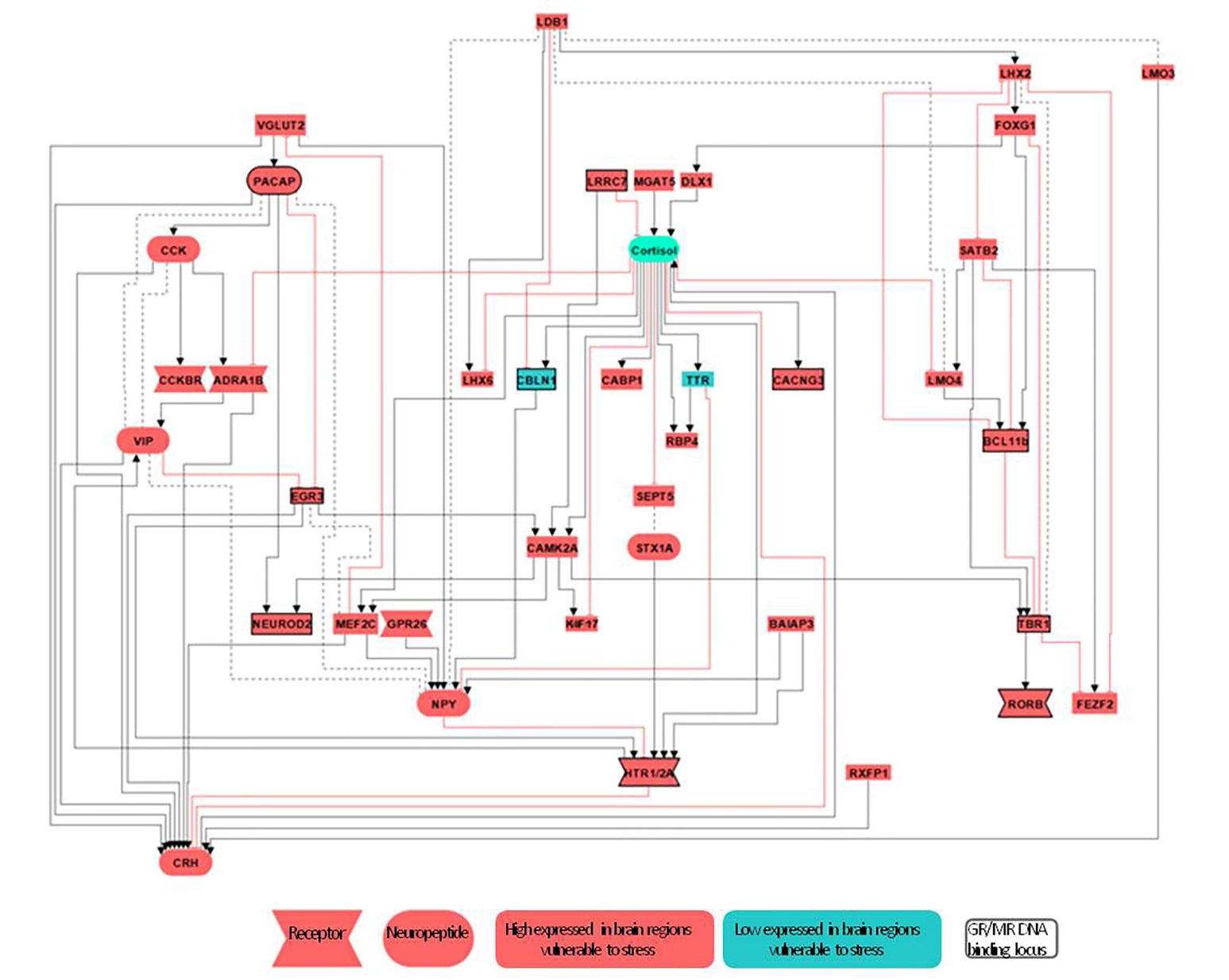

**Figure S7 Putative signaling pathways as the basis of inter-individual differences in stress reactivity** Based on literature search, we assessed whether genes were known to interact with the HPA-axis and each other, resulting in a possible signaling pathway – although spatial specificity is absent from the figure. Red boxes indicate high expression levels of the differentially expressed genes in the brain regions vulnerable to stress, whereas blue boxes indicate low expression. Rounded boxes reflect neuropeptides and notched boxes reflect receptors. Squared boxes indicate the presence of a GR or MR DNA-binding loci in the vicinity of the gene. Black arrows reflect stimulation, the red t-bar indicates inhibition and the dashed line reflects binding or interaction of proteins.
